# Conserved protein folds underpin the diversification of secreted proteins in a fungal pathogen

**DOI:** 10.64898/2026.02.23.707365

**Authors:** Thaís C. S. Dal’Sasso, Eva H. Stukenbrock

## Abstract

**BACKGROUND:** During host colonization, fungal plant pathogens secrete effector-like proteins that alter host cell physiology and target plant-associated microbes. However, rapid evolution and low sequence conservation hinder the study and characterization of these proteins. The fungus *Zymoseptoria passerinii* infects *Hordeum* spp. and includes lineages adapted to wild and domesticated barley. To date, the evolution of effector-like proteins in this species has not been addressed.

**RESULTS:** We combined a set of protein structure-based and network analyses to unravel the secretome of *Z. passerinii*. We first compared AlphaFold2 and ESMFold predictions to establish the baseline for structural analyses. We identified 72 structural clusters in the secretome, revealing fold-level relationships across divergent sequences. We showed that effector-like proteins with host putative immune-interfering functions evolved from a limited group of protein folds, whereas proteins with predicted antimicrobial properties were distributed across fold groups. Physicochemical comparisons indicate that antimicrobial effectors predominantly emerged through amino acid replacements on common effector-enriched scaffolds in *Z. passerinii*, reconfiguring surface charge and electrostatics. We analyzed intra- and interspecific structural variation in selected effector-enriched families by comparing *Z. passerinii* proteins and homologs in the sister species *Z. tritici*. We describe high structural similarity in core folds, with local variation in loop and surface-exposed regions, consistent with fold stability still enabling functional diversification.

**CONCLUSIONS:** The secretome of *Z. passerinii* is organized around common structural folds that support diverse biological roles, including host manipulation and host-associated microbial interactions. Conserved scaffolds combined with local physicochemical variation likely contribute to rapid adaptive evolution in *Z. passerinii*.

## BACKGROUND

Eukaryotic microorganisms use secreted proteins as molecular mediators of their interactions with the environment. Microbial secretomes can participate in nutrient acquisition through enzymatic degradation of substrates, modulate biotic and abiotic stress responses, promote interactions with other organisms, and contribute to growth in diverse habitats [1–4]. Secretome repertoires of pathogens frequently include rapidly evolving proteins that adapt in response to selection pressures imposed by hosts and associated microbes [4]. As a result, secretomes are shaped by gene family dynamics, duplication events, and adaptive evolution linked to ecological changes and niche specialization [5,6]. Thus, secretome repertoires can be surveyed to study microbial adaptation, however extensive genetic variation and low sequence conservation in a majority of secreted fungal proteins challenge the inference of homologies and functions.

In plant-associated fungi, effectors are secreted proteins that play important roles in pathogenesis as they can interfere with host defenses at multiple levels, shaping the outcome of host infection [1,7,8]. Hereafter, we refer to predicted effector candidates as effector-like proteins. Across pathosystems, fungal effectors can alter host cell by suppressing pattern-triggered immunity [9,10], perturb reactive oxygen species signaling [11], alter hormone balance [12], or interact with host components to evade recognition [13,14]. More recent works have documented that some fungal effectors have antimicrobial properties [15]. Such antimicrobial effectors are increasingly recognized as components of plant infection biology as they can inhibit microbial competitors in host tissues or modulate the recruitment of host-associated microbes, thereby reshaping the host niche in favor of the pathogen [3,15,16]. Thus, antimicrobial effectors expand the functional space of effector-like proteins from host-targeting virulence factors to multifunctional molecules that act at the host–pathogen–microbe interface.

Despite their roles in virulence, most fungal effector-like proteins remain uncharacterized due to small size, rapid evolution, and lack of conserved sequence domains [4,17]. For that reason, conventional sequence-based methods often fail to predict their functions as well as their evolutionary relationships among distantly related taxa [18,19], limiting our understanding of their roles in host–microbe interactions. Nevertheless, many effector-like proteins can adopt analogous three-dimensional structures that are important for pathogenesis [6,18,20–22], even in the absence of detectable sequence similarity. These sequence-unrelated but structurally similar effectors [6] exhibit common folds associated with key biological roles during host infection [22]. Therefore, structure-based approaches can be used to uncover protein homologies and similarities not recognized from sequence comparisons, and thereby shed light on mechanisms of fungal pathogenicity that would otherwise remain cryptic among dynamic and fast evolving proteins [6,18,22,23].

The latest developments in protein structure prediction methods provide new opportunities for investigating the effector biology of fungal plant pathogens [22]. Tools such as AlphaFold [24] and ESMFold [25] have allowed the prediction of protein structures on a large scale across a wide range of organisms. These include the plant pathogenic fungi *Magnaporthe oryzae* [20], *Fusarium oxysporum* [26], *Ustilago maydis* [27], and *Venturia inaequalis* [28]. In this context, comparative structural genomics can be used to expand our understanding of effector evolution [6,20]. Effectors families share common ancestry evidenced by conserved protein folds [6]. These highly variable proteins are thought to have diverged through multiple duplication events followed by sequence divergence, sometimes resulting in neofunctionalization and compatibility with new target proteins. [6,22]. However, functional diversification and evolutionary changes at the structural level have so far been poorly explored for secreted proteins of pathogenic fungi.

The fungal genus *Zymoseptoria* includes several grass-infecting species [29]. The most studied species in the genus is the wheat pathogen *Z. tritici*, the causal agent of the Septoria tritici blotch disease [30,31]. A close relative of *Z. tritici, Zymoseptoria passerinii* is the causal agent of Septoria speckled leaf blotch in barley [32]. Although *Z. passerinii* is relatively rare in agricultural production systems, the pathogen persists in natural ecosystems in the Middle East, where it infects species of the genus *Hordeum* [33,34]. These populations of *Z. passerinii* overlap with the center of origin and diversity for Septoria pathogens and their grass hosts [35]. Notably, *Z. passerinii* comprises genetically distinct lineages that infect either wild or domesticated barley, suggesting a long co-evolutionary relationship with the grass hosts [34]. Previously, we characterized the putative effector repertoire in a *Z. passerinii* isolate associated with cultivated barley and further uncovered a large set of unique effectors without sequence homology to effectors predicted in other *Zymoseptoria* species [29]. Moreover, we showed that *Z. passerinii* can alter leaf microbiome composition of wild barley and may engage in interactions with resident microbes [36]. Therefore, *Z. passerinii* likely deploy a diverse array of secreted proteins to manipulate host responses and antimicrobial effectors to mediate microbial manipulation during barley colonization. Thus, *Z. passerinii* serves as an interesting model to study ecological interactions and host specialization of fungal pathogens occurring on wild and cultivated plants [34].

Structure-based analyses offer a promising approach to study rapidly evolving proteins across eukaryotic microorganisms. In this study, we investigate the structural landscape of the secretome of a fungal pathogen, using a *Z. passerinii* isolate from the wild barley species *H. murinum* ssp. *Glaucum* [34] as model system. We hypothesize that structural features of fungal secreted proteins are tied to their biological roles in host colonization and microbial interactions, and that proteins sharing similar folds can act as evolutionary scaffolds supporting functional diversification. To establish a robust baseline for the structural analyses, we first compared the results obtained with AlphaFold2 (AF2) and ESMFold. Although both methods are widely used for protein modelling, they differ in computational cost and runtime, and their accuracy has not been systematically compared for fungal secretomes. We next conducted structural annotation and network-based clustering to identify common [25] folds and uncover homology relationships and similarities within the secretome of *Z. passerinii*, and we sought to determine whether specific folds were associated with effector-like or antimicrobial proteins. We compared physicochemical properties to understand the distribution of antimicrobial activity across effector-like proteins and investigated how conserved structural scaffolds may allow functional diversification. Through an integrative approach, we show that the secretome of *Z. passerinii* is organized around a set of protein folds that support diverse biological roles, and that conserved cores combined with physicochemical variation contribute to effector evolution.

## RESULTS

### Comparison of structural predictions from AlphaFold2 and ESMFold

Due to the low sequence conservation and the absence of conserved protein domains, the functional relevance for most fungal secreted proteins is unknown. We therefore, firstly, set out to establish a robust pipeline for predicting the structures of *Z. passerinii* secreted proteins. We predicted a total of 743 secreted proteins in the proteome of *Z. passerinii* Zpa796 (Additional File 1: Table S1) and used these for structure modeling with AF2 [24] and ESMFold [25]. We predicted a total of 670 protein structures using AF2 and 742 using ESMFold, corresponding to more than 90% of the secretome (Additional File 1: Table S2). Out of 73 mature proteins that failed predictions with AF2, 58 were peptides shorter than 100 amino acids (Fig. 1A-B, Additional File 1: Tables S1 and S2). Overall, AF2-predicted structures displayed significantly higher mean pLDDT values (predicted local distance difference test, p-value < 0.001) than those predicted by ESMFold (Fig. 1, Additional File 2: Fig. S1A). The mean pLDDT value for AF2-predicted structures was 78.23 compared to 63.03 for ESMFold-predicted structures (Additional File 2: Fig. S1A). We note that on the 0–100 pLDDT scale, values above 70 indicate confidently modelled proteins, whereas values below 70 reflect lower prediction accuracy [24,25]. Despite the difference, pLDDT values from AF2 and ESMFold were positively correlated (r = 0.79, p-value < 0.01), although non-linear, across the secretome of *Z. passerinii* Zpa796 (Fig. 1C). To further understand the prediction confidences, we analyzed pLDDT values as a function of mature protein length for the secretome of Zpa796 (Fig. 1D). AF2 structures exhibited significantly higher mean pLDDT values (p-value < 0.01) than ESMFold for proteins up to 900 amino acids long. Structure prediction of short mature proteins (up to 100 residues) remains particularly challenging, but in cases where both software tools successfully predicted a structure, AF2 provided higher pLDDT values than ESMFold (Fig. 1D).

**Fig. 1.**
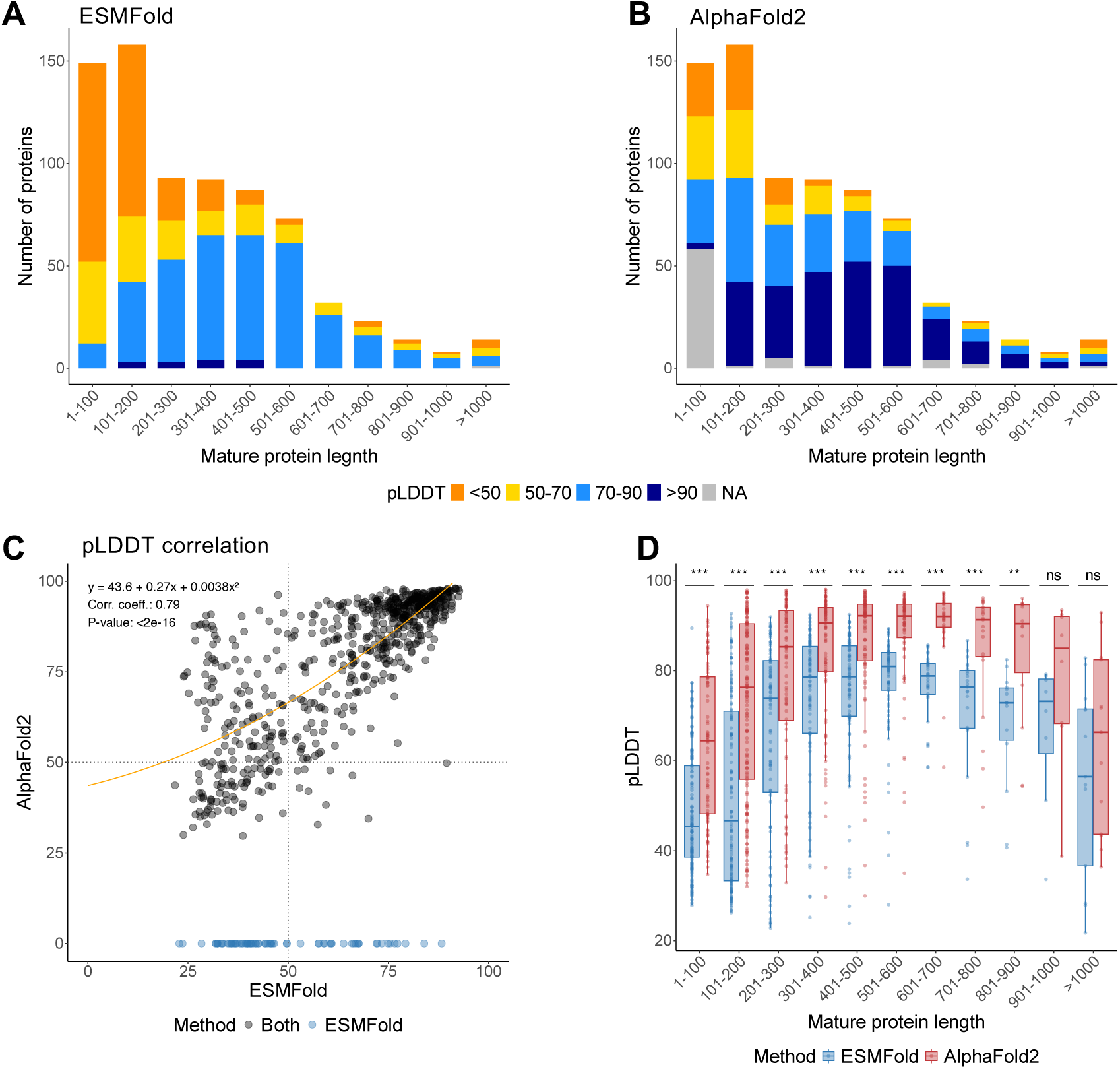
Comparison of protein structural prediction with ESMFold and AlphaFold2 (AF2). Confidence scores of protein structures according to mature protein lengths (number of amino acid residues) shown as the mean per-residue measure of local confidence (pLDDT) values from **A)** ESMFold and **B)** AlphaFold2. Proteins from *Z. passerinii* Zpa796 that failed the structural predictions are labeled as NA (grey). **C)** Spearman’s correlation of pLDDT values for individual protein structures predicted by ESMFold and AF2; proteins uniquely predicted by ESMFold are shown in blue. **D)** Differences in mean pLDDT values per classes of mature protein lengths; significance levels were assessed using the Wilcoxon test (** for p-value <0.01, *** for p-value < 0.001).

We next assessed structural similarity between AF2 and ESMFold predictions for the secretome of *Z. passerinii* Zpa796 using two standard metrics, the template model (TM) score and the root mean square deviation (RMSD), calculated from pairwise alignments of mature proteins predicted by both methods (Additional File 2: Fig. S1B-C, Additional File 1: Table S2). TM-score summarizes global fold similarity and RMSD reports the average distance between aligned Cα atoms, allowing us to test how similar the results from the two prediction methods were. Mature proteins shorter than 100 amino acids had the lowest TM-scores and highest RMSD values between methods, indicating larger structural discrepancies between AF2 and ESMFold models, whereas proteins of intermediate length (401–900 amino acids) showed higher TM-scores and lower RMSD values (Additional File 2: Fig. S1D-F).

To validate whether the results we observed for *Z. passerinii* Zpa796 hold for other fungal proteins, we next benchmarked AF2 and ESMFold predictions against experimentally determined structures using a set of crystallized fungal proteins from the Protein Data Bank (PDB) [37]. In this dataset, AF2 models also showed higher pLDDT values and were more similar to the crystal structures (higher TM-scores, lower RMSD values) than ESMFold models (Additional File 2: Fig. S2, Additional File 1: Table S3). These results are further summarized as Supplementary Text in Additional File 3.

In summary, we conclude that AF2 outperformed ESMFold in terms of prediction confidence and structural accuracy for the fungal proteins which we analyzed here. We therefore used the 670 AF2-predicted structures for *Z. passerinii* for our further downstream analyses.

### Structure-based annotation expands functional coverage of the secretome

We annotated the secretome of *Z. passerinii* Zpa796 using amino acid sequences and AF2-predicted structures (Additional File 2: Fig. S3A, Additional File 1: Tables S1 and S4). Overall, the number of secreted proteins annotated by sequence-based methods was lower than structure-based approaches (Additional File 2: Figs. S3A-C), showing that the structure-based annotation greatly improve our ability to predict functions across the fungal secretome. Of the 670 AF2-predicted structures, 479 (71.5%) were assigned to known structural families using at least one of six reference structural databases (Fig. 2A, Additional File 1: Table S4). We note that proteins of intermediate length (101–600 amino acids) more often matched entries in the protein structural databases, consistent with their higher pLDDT confidence values in this size range (Fig. 2B-D, Additional File 2: Figs. S4-S5).

**Fig. 2.**
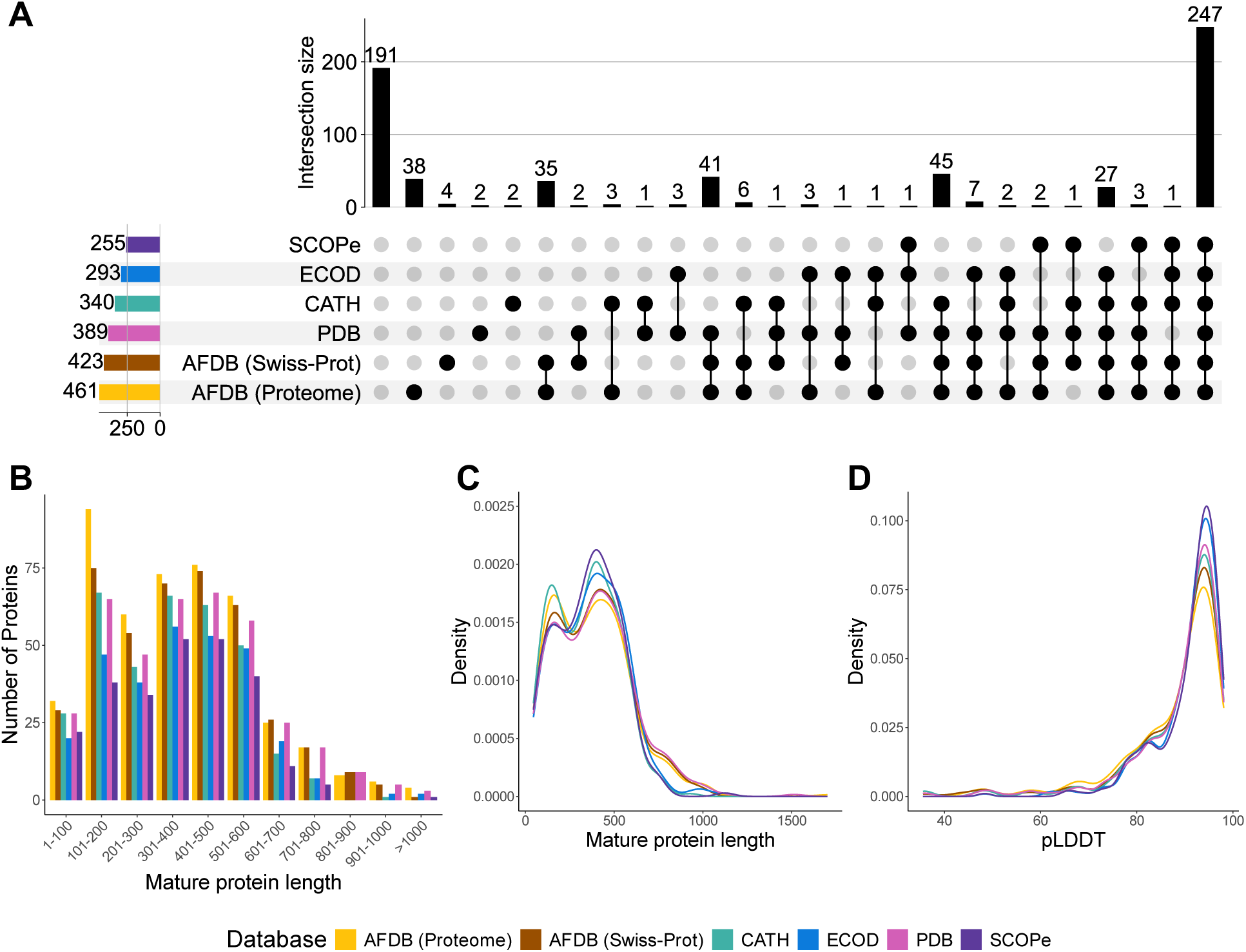
Structure-based annotation of 670 proteins from *Zymoseptoria passerinii* Zpa796. **A)** Upset plot displaying the overlap of protein annotation across structural databases. **B)** Number of proteins annotated by database according to categories of mature protein length. Distribution of protein structures annotated according to **C)** Mature protein length and **D)** pLDDT values for structure databases; AlphaFoldDB (AFDB) subsets “Proteome” and “Swiss-Prot”, CATH, ECOD, PDB and SCOPe.

Among the secreted proteins, we predicted 310 effectors and 341 antimicrobial proteins (AMPs) (Additional File 2: Fig. S3B-C, Additional File 1: Table S1), using the machine learning pipelines EffectorP [19] and AMAPEC [38], respectively. Interestingly, a set of 141 proteins was predicted as both effectors and AMPs, that is, having both immune-suppressing and antimicrobial properties (Additional File 2: Fig. S3D, Additional File 1: Table S1); hereafter, we refer to these proteins as putative antimicrobial effectors.

Within our putative effector and predicted AMP sets, we could predict a function for 143 and 273 proteins, respectively, using protein structures (in comparison using a sequence-based annotation approach we were only able to predict functions for 75 effectors and 220 AMPs) (Additional File 2: Fig. S6). Thus, structure-based data increased functional annotation of effector-like proteins in *Z. passerinii*. A detailed description of the structure-and sequence-based annotations and their comparative overlap is provided in Additional File 3. Per-protein annotations from both sequence- and structure-based methods are listed in Additional File 1: Tables S1 and S4, respectively.

### Structural similarity network

To further explore structural relationships among secreted proteins in *Z. passerinii* Zpa796, we modelled a structural similarity network using the AF2-predicted structures (Fig. 3A). We considered two proteins similar, when bidirectional TM-scores were > 0.5. The network consisted of 364 nodes, each representing a predicted secreted protein structure (Table 1), and 876 edges connecting these proteins according to their similarity. The network displayed a modular structure, with 72 connected components (subgraphs), each representing a structural cluster of proteins showing high structural similarity (Fig. 3A, Table 1, Additional File 1: Table S4). The largest structural cluster comprised 36 proteins (nodes) and the smallest clusters comprised two proteins (26 structural clusters) (Table 1). From the 670 AF2-predicted structures used as input for the network analysis, 306 proteins that did not group into any structural cluster were considered as “singletons” (Additional File 1: Table S4), that is, they lacked detectable structural similarity to other proteins in the Zpa796 secretome. The absence of clustering for singletons reflects a high degree of structural divergence relative to the rest of the secretome. Overall, the network topology reflects a modular secretome architecture in *Z. passerinii* Zpa796, with common folds forming structurally similar groups, and the singletons correspond to the secreted proteins that are structurally unique. Network statistics are described in Additional File 3.

**Fig. 3.**
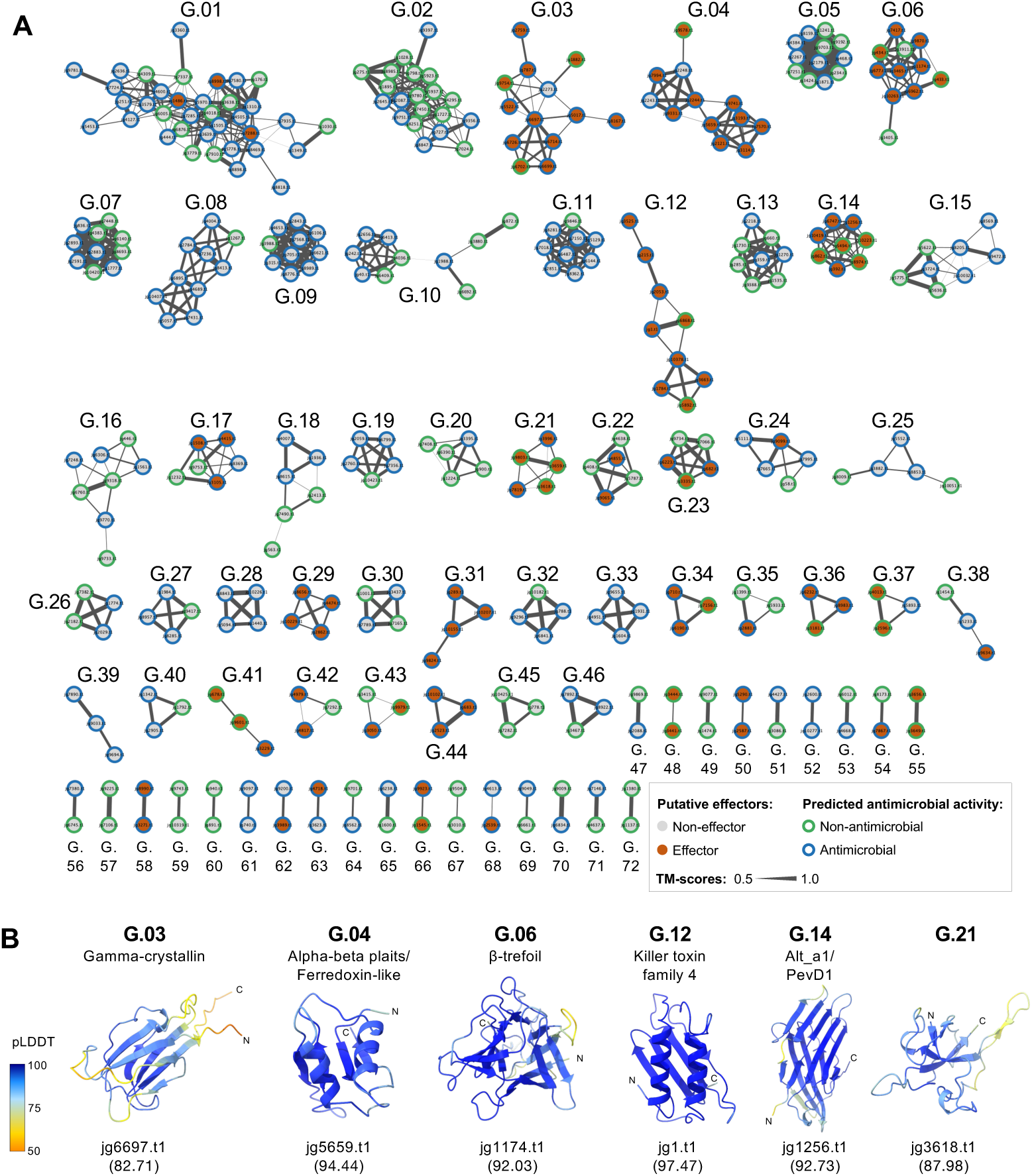
Overview of 72 structural clusters of the *Z. passerinii* Zpa796 secretome. **A)** Graph view of the structural similarity network. The network consists of 72 subgraphs (namely G.01-G.72), each representing a structural cluster. Nodes represent individual proteins and are color-coded according to effector prediction and putative antimicrobial activity. Edges correspond to the structural similarity (TM-score) between pairs of proteins, with thicker edges indicating greater structural similarity. A bidirectional threshold of TM-score > 0.5 was applied to cluster two proteins as structurally similar. **B)** Representative protein structures for each effector-enriched cluster (p-value_adj_ < 0.05). The representative structure corresponds to the node with the highest degree centrality. In cases where multiple proteins had the same centrality value, the representative was selected based on the structure with the highest mean pLDDT value (in parenthesis). Letter codes: N, N-terminus; C, C-terminus.

**Table 1.**
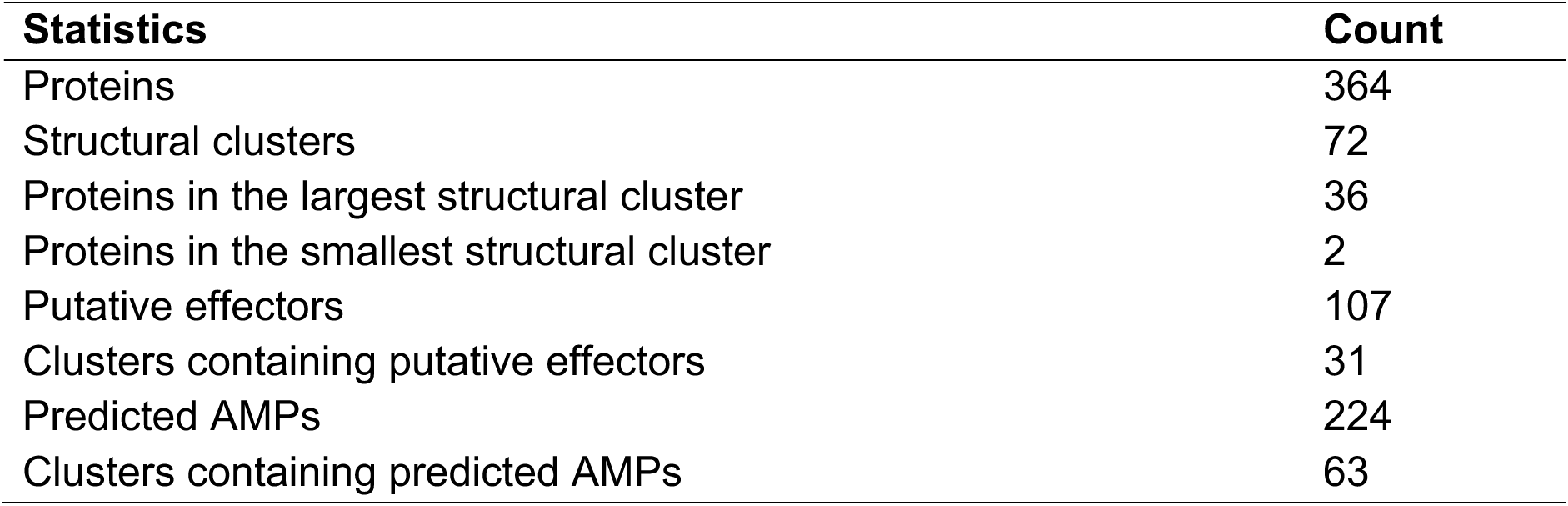
Overview of the proteins and clusters in the structural similarity network.

| Statistics | Count |
| --- | --- |
| Proteins | 364 |
| Structural clusters | 72 |
| Proteins in the largest structural cluster | 36 |
| Proteins in the smallest structural cluster | 2 |
| Putative effectors | 107 |
| Clusters containing putative effectors | 31 |
| Predicted AMPs | 224 |
| Clusters containing predicted AMPs | 63 |

### Functional landscape of structurally similar clusters and structural singletons

We investigated the putative functional context of each structural cluster by integrating our functional annotations (structure- and sequence-based) into the network. Overall, structural clusters were consistent with fold-level annotations from the hierarchical structural databases we tested (CATH, ECOD, and SCOPe). When proteins in a given cluster were annotated using structures, they usually mapped to the same hierarchical category: ‘topology’ in CATH, ‘fold’ in SCOPe, and ‘X-group’ in ECOD (Additional File 1: Table S4). Because these categories reflect similarities in secondary structure and folding patterns [39–42], this agreement indicates that each cluster represents a conserved structural group and provides fold-based, putative functional context for cluster members. Accordingly, clusters in the network spanned diverse protein families, including enzyme-associated clusters (e.g., peptidases, lipases, hydrolases, and oxidoreductases), transport- or binding-related proteins (e.g., immunoglobulin-like and LRR-like domains), fungal surface-associated families (e.g., hydrophobins), toxin-like proteins (e.g., killer toxin proteins), among others (Additional File 1: Table S4). Nevertheless, five clusters (G.21, G.29, G.48, G.55, and G.58) were composed exclusively of uncharacterized proteins (Additional File 1: Table S4). These clusters represent proteins that lack close structural homologs in current databases and may represent fungal-specific proteins.

Putative effectors and predicted AMPs were distributed across the network (Fig. 3A). A total of 107 putative effectors and 224 predicted AMPs were distributed across 31 and 63 structural clusters, respectively (Fig. 3A, Table 1), indicating that diverse folds are used to manipulate plant immunity and to interact with other microbiome members. Among these, 79 proteins were predicted as both effectors and AMPs, that is, putative antimicrobial effectors. Given this distribution, we next tested whether specific structural clusters were enriched, that is, had a higher-than-expected proportion of putative effectors or predicted AMPs (Additional File 1: Table S5). We identified six structural clusters enriched with putative effectors (p-value_adj_ < 0.05) (Fig. 3B, Additional File 1: Table S5): G.03 (Gamma-crystallin proteins), G.04 (Alpha-beta plaits/Ferredoxin-like), G.06 (β-trefoil), G.12 (Killer toxin family 4, KP4), G.14 (Allergen Alt a1, Alt_a1), and G.21 (uncharacterized proteins) (Fig. 3B, Additional File 1: Table S4). No structural cluster was significantly enriched with AMPs (p-value_adj_ > 0.05) (Additional File 1: Table S5) and predicted AMPs were broadly distributed across the different protein folds in the structural network (Fig. 3A). However, clusters with the highest proportion of proteins exclusively predicted as AMPs were usually associated with enzymatic activities (Odds ratios > 2 for AMPs; Additional File 1: Table S5). These include G.08 (Peptidase S8), G.09 (Chloroperoxidases), G.11 (Acid proteases), and G.28 (Peptidase S28), among others (Additional File 1: Table S4). Interestingly, three of the effector-enriched clusters (G.03, G.04, and G.12) also showed a high proportion of putative antimicrobial effectors despite no AMP enrichment (Odds ratios > 2 for AMPs, Additional File 1: Table S5).

Among structural singletons, 218 (71%) lacked structural annotation in the three hierarchical structure databases (CATH, SCOPe, and ECOD) (Additional File 1: Table S4), suggesting they may represent novel folds or adopt conformations not well characterized. Among the remaining 88 singletons, some of the assigned folds included Rossmann-like, immunoglobulin-like, phosphorylase/hydrolase-like, and β-propeller folds (Additional File 1: Table S4). A total of 62 putative antimicrobial effectors were included among the structural singletons.

In summary, we found that putative effectors and predicted AMPs from *Z. passerinii* Zpa796 are distributed across diverse protein folds. Effector-enriched clusters include toxin-like and interaction-associated proteins, consistent with a role of these proteins in host immune interference; meanwhile, clusters with the highest proportions of AMPs were mainly linked to enzymatic activities. A functional overview of other structural clusters is provided in Additional File 3.

### Physicochemical properties across proteins in the network

Since predicted AMPs were dispersed across the structural similarity network, we asked whether predicted antimicrobial activity of *Z. passerinii* proteins is better explained by physicochemical properties than by fold similarity. We evaluated whether a defined set of physicochemical properties distinguished putative antimicrobial effectors from proteins predicted only as effectors or AMPs. We compiled six properties: mature protein length, net charge (pH 7), charge density (net charge per residue), maximum helical hydrophobic moment, mean hydrophobicity, and surface hydrophobicity. We focused on these properties as they capture key biophysical features that contribute to the activity of small secreted proteins, including electrostatic interactions with charged surfaces, membrane association, and protein–protein interactions [43].

To assess if predicted antimicrobial proteins and effectors would differ according to their physiochemical properties, we first grouped proteins from the network into four categories: putative antimicrobial effectors (N=79), proteins predicted solely as putative effectors (N=28), proteins predicted only as AMPs (N=145), and other secreted proteins (N=112). Intriguingly, the four functional groups exhibited significant differences (p-value < 0.05, Kruskal-Wallis test) across all six properties (Fig. 4, Additional File 1: Table S6). Thus, we tested pairwise differences among groups using Dunn’s test. The putative effector-like proteins (predicted as antimicrobial effectors or effectors only) in the network were the shortest and had higher absolute net charge than proteins predicted only as AMPs and other secreted proteins (p-value_adj_ < 0.05) (Fig. 4A-B, Additional File 1: Tables S7 and S8), consistent with small size and higher electrostatic attraction of the effector-like proteins. The net charge values inflated however with increasing protein length (Additional File 2: Fig. S7); thus, we also calculated the charge density (net charge per number of residues) to correct for length differences and enable a more comparable measure of the charge across proteins. Following length normalization, antimicrobial effectors exhibited higher charge density than putative effectors (p-value_adj_ < 0.05), although charge densities overlapped across the other groups (Fig. 4C, Additional File 1: Tables S7 and S8). Interestingly, this pattern is consisted with a shift in amino acid composition toward cationic residues in putative antimicrobial effectors relative to proteins predicted solely as effectors. We also compared different measures of hydrophobicity. The surface hydrophobicity was lower in proteins predicted solely as AMPs, whereas hydrophobic moment and mean hydrophobicity were higher (p-value_adj_ < 0.05) (Fig. 4D–F, Additional File 1: Tables S7 and S8), in agreement with a greater membrane affinity via amphipathic segregation by predicted AMPs when compared to the set of putative effector-like proteins (antimicrobial effectors and effectors only).

**Fig. 4.**
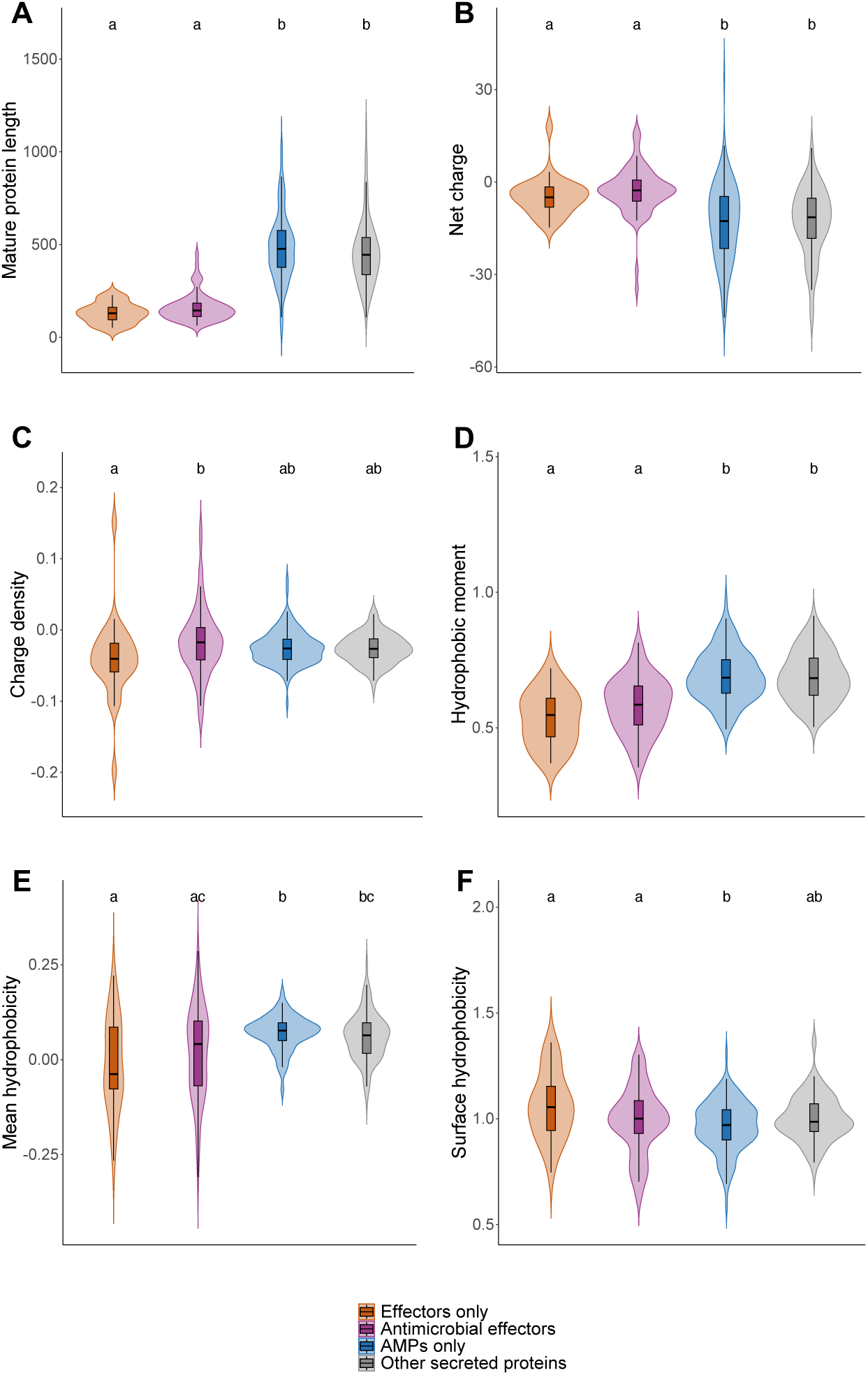
Comparison of physicochemical properties among functional protein groups within the structural similarity network of the *Z. passerinii* Zpa796 secretome. **A)** Mature protein length (number of amino acid residues), **B)** Absolute net charge (pH 7), **C)** Charge density (net charge normalized by the number of residues), **D)** Hydrophobic moment (maximum local helical amphipathicity), **E)** Mean hydrophobicity, and **F)** Surface hydrophobicity. For each property, groups sharing the same letter indicate no significant difference according to Dunn’s test (p-value_adj_ > 0.05). Number of proteins: 28 putative effectors only, 79 putative antimicrobial effectors, 145 predicted only as AMPs, and 112 other secreted proteins.

We repeated the comparisons for each physicochemical property among structurally similar clusters, limiting the analysis to the 46 clusters (63.9% of the 72 total) with at least three protein members (Additional File 2: Fig. S8, Additional File 1: Tables S9-S11). We found significant differences for all six properties in these analyses (p-value < 0.05, Kruskal-Wallis test) (Additional File 1: Table S9). The physicochemical properties that distinguished more pairs of structural clusters were mature protein length, followed by net charge, hydrophobic moment, and mean hydrophobicity (Additional File 2: Fig. S8, Additional File 1: Table S11). Furthermore, several clusters also showed wide within-cluster spreads across the six properties (Additional File 2: Fig. S8), reflecting variation in the physicochemical properties among proteins within the same fold group. Overall, we found significant pairwise differences (p-value_adj_ < 0.05, Dunn’s test) between clusters predominantly composed of effector-like proteins and clusters composed of AMPs or other secreted proteins (Additional File 1: Table S11). We reported the results of physiochemical properties among structural clusters of *Z. passerinii* secretome in Additional File 3.

### High structural similarity and low sequence conservation within clusters

Despite the structural similarity of proteins within clusters, their physicochemical properties varied; we therefore hypothesized that small, localized structural differences among proteins within the same cluster contribute to functional diversification. To explore this, we focused on effector-enriched clusters G.04 and G.12, both containing putative antimicrobial effectors and showing within-cluster variation in physicochemical properties (Fig. 5, Additional File 2: Fig. S8). Overall, we found conserved secondary elements and we observed variation mostly in the loop regions when compared to structured regions (α-helices and β-strands) (Fig. 5).

**Fig. 5.**
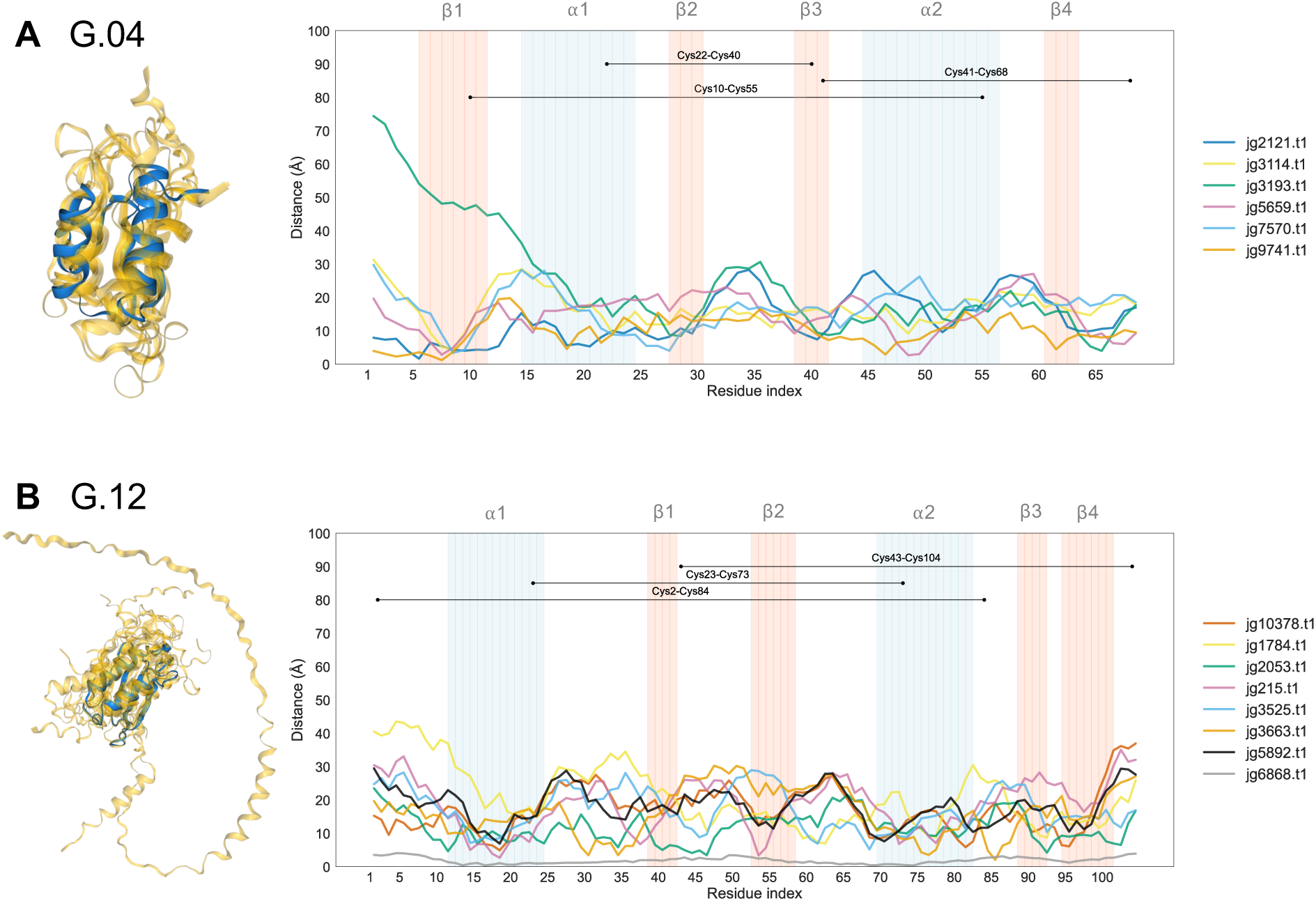
Structural alignments of proteins within structural clusters A) G.04 and B) G.12. On the left, multiple structural alignments with the representative core protein highlighted in blue (jg2244.t1 for G.04, jg1.t1 for G.12, each representative protein corresponding to the shortest protein within the group). On the right, charts showing the distances (in angstroms) between Cα atoms of aligned residues for each cluster member compared to the core protein. Pairs of cysteines forming disulfide bonds and segments forming secondary structures (α-helices in blue and β-strands in red) are highlighted for the core protein of each cluster.

The G.04 cluster comprises proteins annotated as alpha-beta plaits (CATH fold and ECOD X-group) or ferredoxin-like (SCOPe fold), while the G.12 cluster comprises proteins annotated as KP4-like proteins (Fig. 3B, Additional File 1: Table S4). For G.04, protein members varied in structural similarity, so we focused on seven representative proteins that formed the most tightly connected subgroup and were more structurally similar (Fig. 5A). Both G.04 and G.12 clusters consisted of proteins with a α + β organization, meaning they typically contained α-helices and β-strands alternating in a stable scaffold (Fig. 5). The core structure of G.04 aligned proteins (represented by protein jg2244.t1) were organized in a β-α-β-β-α-β arrangement, with five loop regions (3–9 residues) connecting the secondary structure elements (Fig. 5A). In the representative G.04 protein, we predicted six cysteines forming three disulfide bridges: Cys10–Cys55 (5.23 Å between α-carbons, Cα), Cys22–Cys40 (5.93 Å), and Cys41–Cys68 (6.42 Å) (Fig. 5A), which contribute to fold stability. Within G.04, the first (β1) and third (β3) β-strands exhibited the smallest distance variation between aligned amino acids, indicating the most conserved secondary elements among proteins. Meanwhile, the loops β1–α1 and β2–β3 displayed the largest deviations from the core protein fold (jg2244.t1, Fig. 5A). Among G.12 aligned proteins, the core structure (represented by jg1.t1) displayed an α-β-β-α-β-β organization, connected by five loops (2–14 residues) (Fig. 5B). In the representative G.12 protein, we also predicted six cysteines forming three disulfide bridges: Cys2–Cys84 (5.10 Å between Cα), Cys23–Cys73 (5.72 Å), and Cys43–Cys104 (6.40 Å) (Fig. 5B). Among G.12 proteins, the smallest distances between aligned residues were observed within the α-helices (α1 and α2) and the largest in the loop α1–β1, when compared to the core protein fold (jg1.t1, Fig. 5B). Taken together, both clusters retain a conserved, disulfide-stabilized core fold, while most structural differences concentrate in surface-exposed loop regions.

Despite structural similarity, we found low sequence conservation within G.04 and G.12 clusters (Additional File 2: Figs. S9-S10). In G.04, four cysteines were conserved across all members, whereas most other positions varied and included conservative and non-conservative replacements (Additional File 2: Fig. S9A). Meanwhile in G.12, sequence conservation was even lower, with only one cysteine and one tryptophan conserved across all sequences (Additional File 2: Fig. S9B). Overall, the sequence alignment patterns mirrored the structural clusters identified through network analysis, yet structural similarity exceeded sequence identity between pairs of proteins (Additional File 2: Fig. S10). Even pairs of proteins with nearly identical structures displayed several non-conservative amino acid replacements that result in low sequence identity, for example paralogs jg1.t1 (putative antimicrobial effector) and jg6868.t1 (putative effector only) (RMSD = 0.46 and TM-score = 0.986, normalized for both proteins) from G.12 cluster (Additional File 2: Figs. S9B and S10B).

In summary, proteins within clusters G.04 and G.12 retained core folds despite low sequence conservation. Variation arose from amino acid replacements that contribute to differences in protein charges and loops varying in length, consistent with these stable cores serving as scaffolds that allow the evolution of different protein functions while preserving a conserved fold.

### Evolution of protein structures between homologs of *Z. passerinii* and *Z. tritici*

To gain insights into the evolution of the structurally similar proteins across species, we searched for homologs within the *Zymoseptoria* genus. We used OrthoFinder to identify homologous relationships between secreted proteins of *Z. passerinii* Zpa796 and other *Zymoseptoria* species. Of the 364 proteins in the structural similarity network, 359 had homologs in at least one other species (Additional File 1: Table S12). Overall, most proteins from Zpa796 within a given structural cluster fell into different orthogroups along with homologs from the other species (Additional File 1: Tables S4 and S12), indicating that their primary sequences have diverged greatly both within *Z. passerinii* and across close related species.

We selected the proteins jg9741.t1 and jg3663.t1 (cluster G.04 and G.12, respectively) for detailed structural comparison with their homologs in the reference isolate *Z. tritici* IPO323 (Fig. 6). Protein jg3663.t1 had a single ortholog in IPO323 (ZtIPO323_026410.1), while jg9741.t1 had two paralogs: ZtIPO323_014270.1 and ZtIPO323_104450.1 (Additional File 1: Table S12). See Additional File 1: Table S13 for aliases of these IPO323 proteins across annotation versions. Pairwise structural alignments of homologs revealed highly similar folds, with small differences found in surface-exposed loops and terminal extensions (Fig. 6), supporting overall fold conservation of these proteins across the sister species. Additional information on homologs distribution and alignment scores are provided in Additional File 3.

**Fig. 6.**
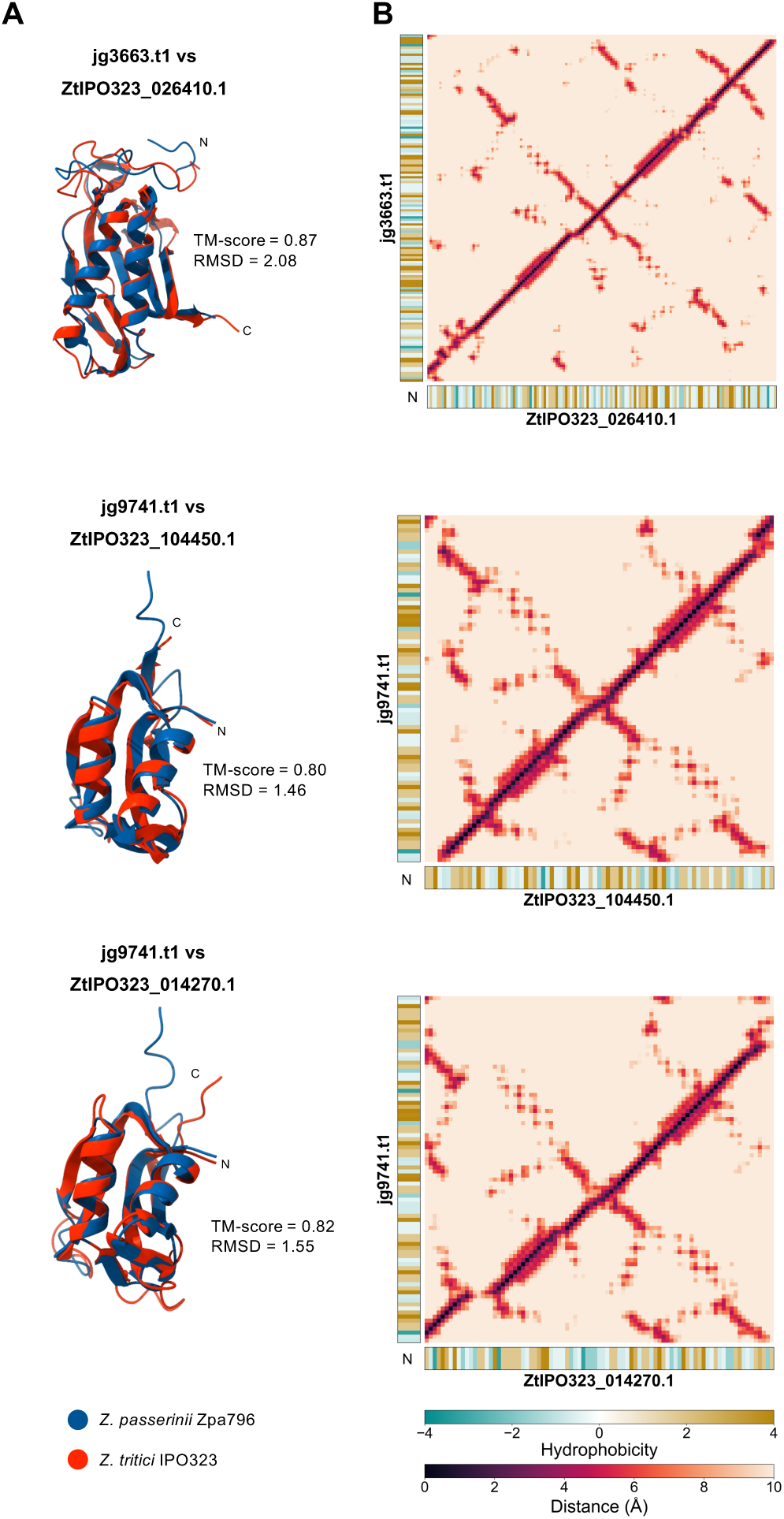
Structural comparison between *Z. passerinii* Zpa796 proteins and their homologs in *Z. tritici* IPO323. The alpha-beta plait (jg3663.t1) and KP4-like (jg9741.t1) proteins from *Z. passerinii* Zpa796 belong to effector-enriched structural clusters G.04 and G.12, respectively. **A)** Superposition of tertiary structures, with TM-scores normalized to the *Z. passerinii* Zpa796 proteins (shown in blue). **B)** Symmetric distance matrix comparing all pairs of amino acid residues, measured using Cα atoms as reference. Helices are represented as strips directly adjacent to the diagonal, while antiparallel β-strands appear as cross-diagonal patterns. Hydrophobicity per residue was calculated using the Kyte-Doolittle scale. Proteins from *Z. passerinii* Zpa796 are displayed on the y-axis, and proteins from *Z. tritici* IPO323 (according to [58]) on the x-axis. Letter N indicate N-termini.

We next assessed whether sequence divergence between species leads to differences in protein properties. We compared physicochemical properties associated with antimicrobial activity for jg3663.t1 and jg9741.t1 and their homologs in *Z. tritici* IPO323 (Additional File 1: Table S14). Both Zpa796 proteins and their IPO323 homologs were annotated as antimicrobial effectors. The main differences among homologs were in net charge and isoelectric point (pI), with the *Z. passerinii* Zpa796 proteins exhibiting higher net charges and pI values than their counterparts in *Z. tritici* IPO323 (Additional File 1: Table S14). Accordingly, the homologs from Zpa796 should remain more positively charged at mildly acidic host apoplast than the homologs from IPO323. Although hydrophobic moment was comparable across homologs of the two species (Additional File 1: Table S14), jg9741.t1 (G.04) and ZtIPO323_014270.1 displayed a more hydrophobic nonpolar face for helix α1 (amphipathic helix) (Fig. 6B, Additional File 1: Fig. S11) and higher surface hydrophobicity than the IPO323 paralog ZtIPO323_104450.1 (Additional File 1: Table S14). Mean hydrophobicity and Boman index were broadly similar across homologs (Additional File 1: Table S14). Together, these data indicate species- and paralog-specific variation in surface properties for these proteins with conserved structural scaffold.

## DISCUSSION

We explored the structural landscape of secreted proteins in the wild grass pathogen *Z. passerinii* using an array of protein structure analyses. By predicting and clustering proteins by structural similarity, we describe common folds in sequence-divergent proteins that may engage in host manipulation and plant-associated microbial interactions. An interesting finding is that a large proportion (>45%) of all secreted proteins are predicted to encode antimicrobial proteins pointing to an un-acknowledged importance of microbial interactions during the life cycle of *Z. passerinii*. Furthermore, we find that many effector proteins also are predicted to be antimicrobial. This suggest that the same protein folds, perhaps even the same proteins, can be used for highly distinct purposes in the host tissue.

To explore protein functions and evolution in the secretome of *Z. passerinii* we started our investigations with a detailed assessment and comparison of predicted protein structures. We first evaluated the structural predictions generated by AF2 and ESMFold and obtained models with higher confidence values for AF2 predictions, particularly for proteins with conserved domains. In contrast, predictions for short proteins were more variable, with AF2 leaving some sequences unmodeled and ESMFold assigning very low-confidence models. These unmodeled sequences were frequently predicted as intrinsically disordered, consistent with more dynamic conformational behavior [44] in this set of proteins.

Protein annotation and network-based clustering of AF2-predicted structures revealed the relationships across proteins of the secretome of *Z. passerinii* Zpa796. The fragmented topology of the structural similarity network reflected a broad structural diversity among the secreted proteins. Instead of forming a densely connected architecture, structurally similar proteins grouped into discrete clusters corresponding to protein families associated with distinct biological functions. Among the effector-enriched clusters, were two structural families, Alt a1–like (G.14) and β-trefoil (G.06), containing folds previously implicated in host immune modulation by fungal plant pathogens [45,46]. Alt a1–like proteins can interfere with ROS signaling [47], inhibit antifungal pathogenesis-related proteins [48], and show induced gene expression in *Zymoseptoria* species during host colonization [49]. Meanwhile, β-trefoil proteins are commonly associated with fungal lectins and non-enzymatic protease inhibitors involved in diverse ligand interactions [50–52].

Other effector-enriched folds in the secretome of *Z. passerinii* Zpa796 correspond to proteins with the potential to engage both in host defenses and microbial interactions. These folds encompassed KP4-like (G.12) [53–55], alpha-beta plait (ferredoxin-like; G.04) [56,57], and gamma-crystallin-like (G.03) [57] proteins. Crystallin-like effectors are widespread across fungi [6,20,21], and usually display induced gene expression during early host infection [57]. Meanwhile, KP4-like effectors contribute to ion channel interference, as shown in *U. maydis* [55], and PAMP-triggered immunity (PTI) suppression, as shown in *Z. tritici* [58] and *F. graminearum* [53]. Similarly, alpha-beta plaits, which include some variants as the killer toxin family 6 (KP6)-like subfamily, exhibit functional divergence across fungi, being able to modulate effector-triggered immunity [58] or inducing host cell death [57]. Moreover, both KP4-like and KP6-like variants of alpha-beta plaits from *Z. tritici* may have antifungal activity *in vitro* [59]. Therefore, the plethora of putative antimicrobial effectors within these effector-enriched clusters from *Z. passerinii* suggests that these fungal proteins have evolved multifunctionality in plant-associated niches [15,60].

Interestingly, some effector-associated clusters remained structurally unclassified despite structural similarity among members (i.e. G.21, effector-enriched). In these cases, proteins showed a prevalence of intrinsically disordered regions that contribute to structural flexibility and interactions with multiple molecular targets, which can lead to neo-functionalization of some proteins [61]. It is plausible to consider that the abundance of disordered proteins in the secretome of *Z. passerinii* Zpa796 also contributes to the functional versatility and evolutionary diversification of proteins involved in host–microbe interactions.

Unlike putative effector proteins, predicted AMPs did not achieve fold-level enrichment, suggesting that antimicrobial activity depends on physicochemical properties recurring across diverse protein folds. A key discriminator between proteins predicted exclusively as AMPs and putative effector-like proteins was helical amphipathic patterning, measured by the hydrophobic moment. Amphipathic α-helices are associated with antimicrobial activity due to their ability to interact with and damage microbial membranes [43,62–64]. In *Z. passerinii* Zpa796, greater hydrophobicity and enhanced amphipathicity among proteins predicted exclusively as AMPs indicate a mechanism in which helix-face segregation facilitates microbial membrane insertion and disruption [64] by these proteins. Conversely, effector-like proteins in the network were shorter and showed higher absolute net charge compared to other proteins in the secretome. Notably, putative antimicrobial effectors exhibited greater charge density (net charge per residue), reflecting an enrichment in positively charged residues compared with proteins predicted exclusively as effectors. An increased cationicity among antimicrobial effectors is suited to engaging negatively charged surfaces in the host [65,66] and on microbial membranes [43,64], therefore, enhancing membrane-disruptive potential while retaining an effector conserved protein fold. Mapping these properties across the structural similarity network reinforced this pattern as putative antimicrobial effectors were abundant within effector-enriched clusters, which allowed us to hypothesize how these proteins evolved.

Two complementary mechanisms may contribute to the functional diversification of effector-like proteins in *Z. passerinii* Zpa796 while retaining common protein folds: i) non-conservative amino acid replacements and ii) the maintenance of flexible loop and linker regions.

First, our results are consistent with antimicrobial effectors emerging through amino acid replacements on effector-enriched structural scaffolds, as reflected by their clustering with effector-enriched folds and shared physicochemical profiles. As a result, antimicrobial properties in effector-like proteins may have arisen through amino acid replacements that alter the surface electrostatic potential of effector backbones. Likewise, amino acid substitutions that reconfigure surface properties are recurrent in common effector folds across fungal plant pathogens [21,66–68]. Nevertheless, alternative evolutionary routes are plausible, including antimicrobial effectors evolving from ancestral AMPs that later diversified toward host manipulation [15]. Thus, while our dataset supports effector-scaffold diversification as the dominant pattern in *Z. passerinii* Zpa796, it does not exclude AMP-derived trajectories in specific proteins.

Second, conserved structural cores paired with flexible loops and linker regions contribute to diversification within common effector families. Although G.04 (alpha–beta plait/ferredoxin-like) and G.12 (KP4-like) proteins retained highly similar core folds, we observed most structural variation in loop regions. Variation in loop length and composition can influence ligand binding and molecular recognition [69–71]. Similarly, loop flexibility has been linked to functional shifts in fungal secreted proteins, including cerato-platanins from *Ustilago* spp. [72] and the acquisition of antimicrobial properties by a group of fungal chitinases [73]. In *Z. passerinii* Zpa796, the combination of a stable structural core and flexible loop regions enable these proteins to maintain their overall fold while contributing to new putative functions or allowing different binding partners throughout host infection.

Comparing two putative antimicrobial effectors of *Z. passerinii* Zpa796 with their homologs *Z. tritici* IPO323 shed further light on how structurally similar effectors evolve and diversify across related fungal pathogens. We focused on the Zpa796 proteins jg3663.t1 (a KP4-like protein, G.12) and jg9741.t1 (an alpha–beta plait protein, G.04) for their placement in effector-enriched clusters and predicted antimicrobial activity. Notably, their homologs in IPO323 are effectors that induce plant-immune suppression [58], providing a functional framework for our findings in Zpa796. Despite high structural similarity, differences in surface-exposed regions and amino acid composition among *Z. passerinii* Zpa796 and *Z. tritici* IPO323 homologs may reflect host adaptation. Although the leaf apoplast of grasses is mildly acidic during biotic stress [74,75], the higher isoelectric point (pI) and net charge of Zpa796 proteins relative to their IPO323 counterparts likely reflect host-specific tuning of the surface charge, enhancing interactions with anionic cell-wall components and microbial membranes in the barley apoplast when compared with conditions experienced by *Z. tritici* in wheat. Experimental test in the native host or with plant-specific receptors are necessary to test this hypothesis.

Notably, amphipathic α-helices were an additional characteristic of jg9741.t1 from Z. *passerinii* Zpa796 and its homologs in *Z. tritici* IPO323, all adopting alpha–beta plait folds. IPO323 homologs are KP6 variants within the alpha-beta plaits group [58], in which amphipathic α1 helices contribute to the disruption of microbial membranes [76]. Preliminary confrontation assays showed that homolog ZtIPO323_014270.1 inhibits *in vitro* growth of *Escherichia coli* (Thynne et al., unpublished), further supported by the antimicrobial activity among killer toxin–like effectors from *Z. tritici* [59]. Together, these experimental findings support our *in silico* analyses of antimicrobial effectors in *Z. passerinii* and validate our list of candidates for future experiments aimed at elucidating how these proteins interact with microbial membranes or host components.

## CONCLUSIONS

Using a suite of deep learning tools, we find that the secretome of the fungal pathogen *Z. passerinii* comprises a diverse repertoire of proteins that appear to have evolved from a limited number of common protein folds. Within groups of structurally similar proteins, local variation in surface-exposed regions and physicochemical properties provides a basis for functional diversification, including host-targeting activities and antimicrobial properties. Our findings suggest that a large proportion of proteins secreted by fungal pathogens are produced to interfere with other microorganisms in the same niche. Together, this secretome-wide study provides a valuable source of effector candidates for experimental functional elucidation and a general strategy that can be transferred to the study of secretomes from other eukaryotic microorganisms.

## METHODS

### Assembly of genomic and crystallized protein structure data

We retrieved the genome sequence, gene models, and 10,461 protein sequences of the *Z. passerinii* isolate Zpa796 (obtained from the wild barley species *H. murimum* ssp. *glaucum*), described previously in [34], (Additional File 1: Table S1). From the Protein Data Bank (PDB v240622, [37]), we retrieved a dataset of 228 fungal proteins, consisting of crystallized structures and amino acid sequences (Additional File 1: Table S3). We filtered entries in the PDB using the text terms “fungal effector” and “fungal protein.” Only proteins from fungal organisms within the phyla Ascomycota and Basidiomycota were kept. Structures crystallized with a single chain were retained, and the final dataset was manually curated to reduce protein redundancy.

### Sequence-based protein functional annotation

We performed sequence-based annotation of the *Z. passerinii* Zpa796 proteome to identify conserved protein domains and functional signatures. Protein domains were predicted using InterProScan v5.54-87.0 [77]. EuKaryotic Orthologous Groups (KOG), Gene Ontology, and KEGG annotations were retrieved using eggNOG-mapper v2 [78]. Functional classes such as CAZymes, lipases, and proteases were identified based on hidden Markov model (HMM) or BLASTp searches against specialized databases. Secondary metabolite biosynthesis clusters were predicted using antiSMASH v7.1.0 [79]. Detailed parameters, database versions, and thresholds are provided as Supplementary Text in Additional File 3.

### Secretome and effector prediction

We defined the predicted secretome by the presence of a signal peptide, predicted by SignalP v6.0 [80] and Phobius v1.01 [81], and the absence of transmembrane domains, predicted by TMHMM v2.0 [82] and Phobius. We used TargetP v2.0 [83] and DeepLoc v2.0 [84] to predict the subcellular localization of the secreted proteins. ESpritz v1.3 [85] was used to search for intrinsically disordered regions in the candidate secreted proteins. Fungal effectors are typically soluble secreted proteins that enter the apoplast or are delivered inside host cells [4,19]. We defined the putative effectors of *Z. passerinii* Zpa796 using EffectorP v3.0 [19] set as fungi.

### Structure prediction

Protein structures for the candidate secreted proteins were obtained using the mature sequences, with predicted signal peptides removed prior to modeling. We conducted structural prediction locally using AlphaFold2 (AF2) v2.3.1 [24] with the parameter “--max_template_date=2020-05-01” and ESMFold v1.0.3 [25] with “--chunk-size 32.” For AF2, five models were generated for each protein, and we selected the highest-quality model (ranked_0.pdb) based on the highest average pLDDT (predicted local distance difference test) score.

### Comparison of AlphaFold2 and ESMFold predictions

Fungal secreted proteins are highly variable in size, and many fungal proteins involved in plant–microbe interactions correspond to short amino acid sequences [4,17,19]. We compared structural predictions across the full secretome of *Z. passerinii* Zpa796 and also assessed prediction quality across protein length categories (in 100-residue intervals). For proteins predicted by both methods, we compared structural confidence (pLDDT values) and structural similarity. We used TM-align v20220412 [86] to calculate the template modeling (TM) score and root mean square deviation (RMSD) values. Additionally, we evaluated how similar the outputs from each method were to experimentally determined fungal structures by aligning predicted models to crystallized proteins retrieved from the PDB. Statistical analysis and comparison metrics are described in the Additional File 3.

### Structural annotations and antimicrobial activity prediction

We used the *easy-search* workflow from Foldseek [87] to perform structural similarity searches against the following protein structure databases: CATH50 v4.3.0 [39], SCOPe40 v2.08 [40], ECOD40 v291 [41], PDB v240101 [37] and AlphaFold DB v4 (AFDB) [88] with separate runs for “Proteome” and “Swiss-Prot” entries. The Proteome section of the AFDB contained predicted structures for all proteins in reference proteomes, the Swiss-Prot section included predicted structures from UniProt entries that were manually curated. Hereafter, we refer to these as AFDB (Proteome) and AFDB (Swiss-Prot). Searches were conducted with the parameters “-s 7.5 --alignment-type 1 --tmscore-threshold 0.5 --format-mode 4” and a customized “--format-output” parameter with several output columns, including “query,target,qcov,qtmscore”. We defined the best hit for each *Z. passerinii* Zpa796 protein based on the highest query coverage (qcov) and the highest query TM-score (qtmscore). Custom Python3 scripts were used to extract structural annotations from the best-hit entries in the reference databases. We predicted antimicrobial activity among candidate secreted proteins using the machine-learning pipeline AMAPEC v1.0b [38], with protein structure files generated by AF2 as inputs.

### Modelling and analysis of the structural similarity network

We built a structural similarity network to investigate protein homology relationships within the predicted secretome of *Z. passerinii* Zpa796. Structural similarity between secreted proteins was calculated using all-vs-all pairwise alignment with TM-align. Two structures were considered similar using as cutoff a TM-score > 0.5 normalized for both proteins. We modeled the network using NetworkX [89] in Python3, based on the following criteria: i) candidate secreted proteins were represented as nodes in the graph, with each node corresponding to one protein; ii) connections between nodes were represented by edges when the bidirectional TM-score cutoff was met, the highest TM-score from each pairwise comparison was used as edge weight for plotting. We further analyzed the modeled network by calculating network statistics and metrics using NetworkX. The network was visualized using Cytoscape v3.10.3 [90]. We treated each network subgraph as an individual structural cluster, representing a group of proteins with similar structural features that might reflect functional or evolutionary relationships. To identify significant protein folds involved in *Z. passerinii* Zpa796 interactions with the host and other microorganisms during plant infection, we tested for enrichment of putative effectors and candidate antimicrobial proteins within structural clusters using Fisher’s exact test (p-value_adj_ < 0.05, using the Benjamini–Hochberg correction) from the SciPy v1.13.0 [91] library in Python3.

### Structural comparisons within *Z. passerinii* clusters and *Zymoseptoria* homologs

We investigated sequence conservation among proteins belonging to the same structural cluster by generating multiple sequence alignments (MSAs) using the L-INS-i algorithm implemented in MAFFT v7.525 [92]. Alignments were trimmed with trimAl v1.5.rev0 [93] using the “-gappyout” option and visualized using the *ggmsa* package [94] in R v4.3.3 [95]. Pairwise percent identity was computed from MSA using the Bio.Phylo.TreeConstruction module from Biopython v1.78 [96]. To assess structural conservation, we used FoldMason [97] with default parameters to perform multiple structure alignments of proteins within a structural cluster. To quantify the variation between aligned residues, Euclidean distances were calculated using Cα atoms as reference points, implemented via the Bio.PDB module from Biopython.

Sequence-based homologous relationships among *Z. passerinii* Zpa796 and other *Zymoseptoria* species were identified using OrthoFinder v2.2.7 [98] through an all-versus-all comparison with DIAMOND v0.9.24.125 [99]. Homologous proteins from *Z. passerinii* Zpa796 and *Z. tritici* IPO323 were aligned using TM-align. Protein structures from *Z. tritici* were based on the reference genome of the isolate IPO323 and retrieved from [58]. Distance maps for pairwise structural alignments were generated using Cα atoms to calculate residue-residue distances. For aligned protein pairs, residue-level hydrophobicity was calculated using the Kyte-Doolittle scale [100].

### Comparison of physicochemical properties of proteins

The antimicrobial activity of proteins is also governed by physicochemical properties that influence their interactions with microbial membranes [43]. We quantified four properties from mature protein sequences using AMAPEC: net charge (at pH 7), hydrophobic moment (computed as the maximum helical amphipathicity), mean hydrophobicity, and surface hydrophobicity. Parameters used by AMAPEC to compute each of these four physicochemical properties are described in Additional File 3. Additionally, we calculated charge density, defined as absolute net charge normalized by sequence length (net charge per residue). Statistical analyses were performed on proteins present in the structural similarity network, considering these five physicochemical properties plus mature protein length. The six properties were evaluated at two levels (functional groups of proteins in the network and structural clusters) in R v4.3.3. Differences among functional groups and structural clusters were assessed with Kruskal–Wallis test (p-value < 0.05) followed by Dunn’s test for post hoc pairwise comparisons (p-value_adj_ < 0.05, using the Benjamini–Hochberg correction). Effect sizes were calculated using Cohen’s d. For cluster-level analyses, we restricted comparisons to clusters with at least three proteins to ensure statistical rigor.

## DECLARATIONS

## Data availability

Protein structures, network graph file, and metadata generated in this study are available at Zenodo (https://doi.org/10.5281/zenodo.16792728). Code for the reproduction of the analyses in this paper is available across GitHub repositories: for comparative structural genomics (https://github.com/thaisdalsasso/Structural_genomics_Zpa796) and protein annotation (https://github.com/thaisdalsasso/Protein_annotation).

## Authors contributions

TCSD: Conceptualization, Formal analysis, Investigation, Methodology, Visualization, Data curation, Writing. EHS: Conceptualization, Investigation, Funding acquisition, Project administration, Writing.

## Acknowledgements

We thank Idalia Rojas-Barrera for sharing the genome and gene models of *Z. passerinii* Zpa796 prior to publication. We thank all members of the Environmental Genomics group for their valuable discussions and feedback. We also acknowledge the high-performance computing resources and support provided by the Kiel University Computing Centre, the Max Planck Institute for Evolutionary Biology, and the GWDG Göttingen.

## SUPPORTING INFORMATION

**Additional File 1:** Supplementary Tables S1-S14

**Additional File 2:** Supplementary Figures S1-S11

**Additional File 3:** Supplementary Text

## Abbreviations

AF2: AlphaFold2
AMPs: antimicrobial proteins
pLDDT: predicted local distance difference test
KP4: killer toxin family 4
KP6: killer toxin family 6
PDB: Protein Data Bank
pI: isoelectric point
RMSD: root mean square deviation
TM: template modeling

