## Additional File 2 for "Conserved protein folds underpin the diversification of secreted proteins in a fungal pathogen"

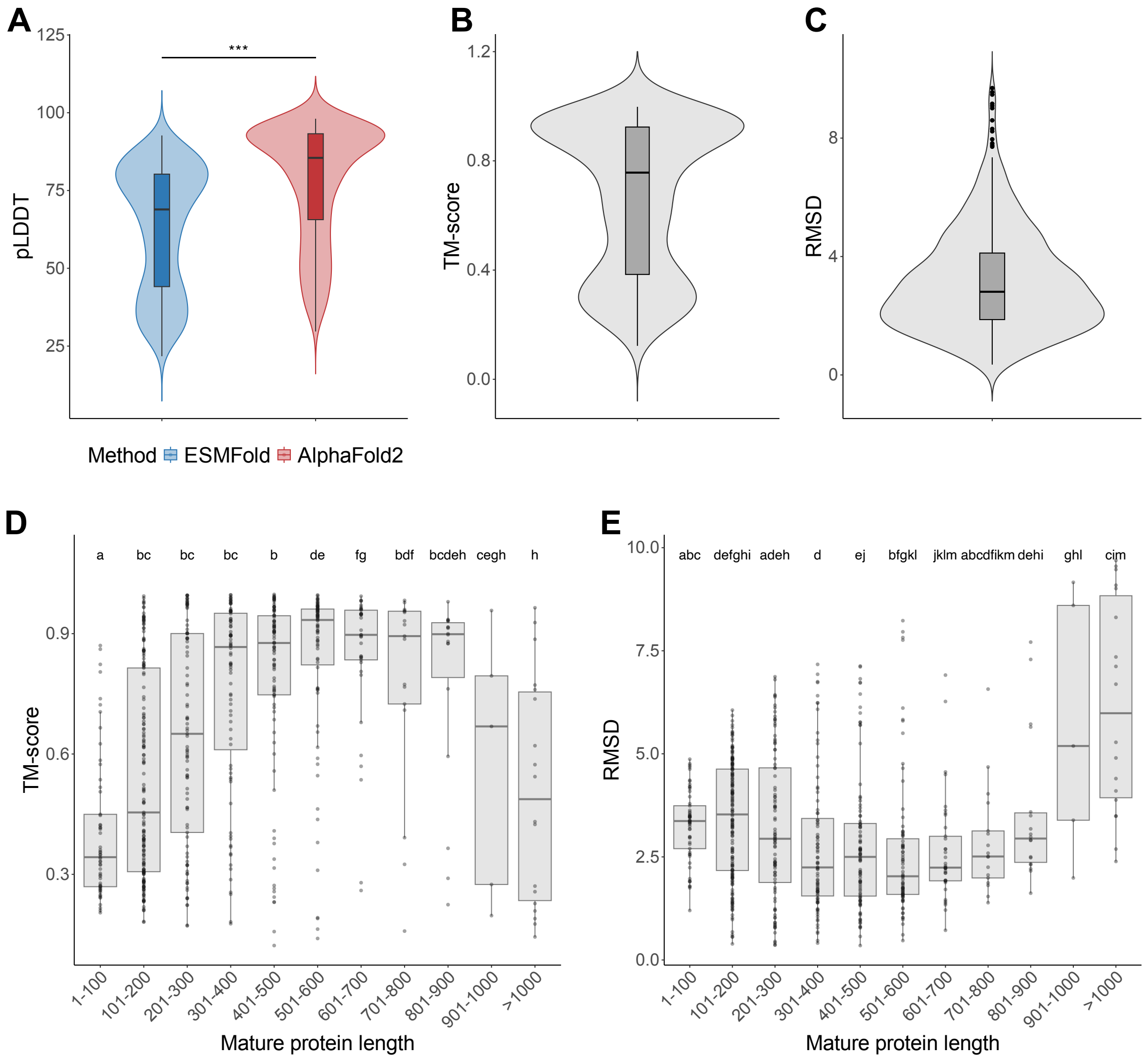


**Fig. S1. Comparison of protein structural prediction using ESMFold and AlphaFold2 (AF2). A)** Difference in mean pLDDT values for all proteins predicted by the two methods; significance differences between methods were assessed using the Wilcoxon test (*** for p-value < 0.001). Mean structural similarity measured by **B)** Template modeling (TM) scores and **C)** Root-mean-square Deviation (RMSD) for all proteins predicted by ESMFold and AF2. Mean structural similarity given as **D)** TM-scores and **E)** RMSD values between the two prediction methods according to mature protein length; length ranges sharing the same letter indicate no significant difference according to Dunn's test ($\text{p-value}_{adj} > 0.05$)

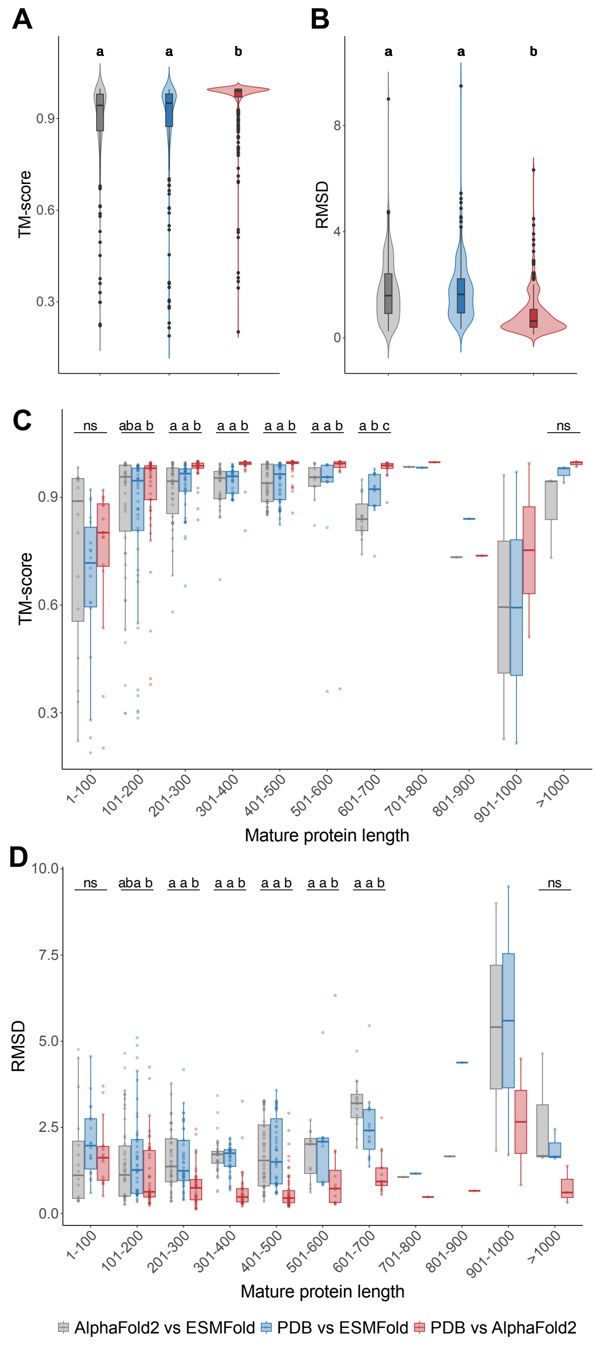


**Fig. S2. Comparison of structural similarities between fungal crystallized protein structures (retrieved from the Protein Data Bank, PDB) and their predicted structures by ESMFold and AF2.** Mean structural similarity measured by **A)** Template modeling (TM) score and **B)** Root-Mean-Square deviation (RMSD). Structural similarity for each protein length range given as **C)** TM-scores and **D)** RMSD values; comparisons sharing the same letter indicate no significant difference according to Dunn's test (p-value$_{adj}$ > 0.05). No statistical test was performed for proteins 701–900 amino acids long due to the number of samples included (fewer than three proteins per category).

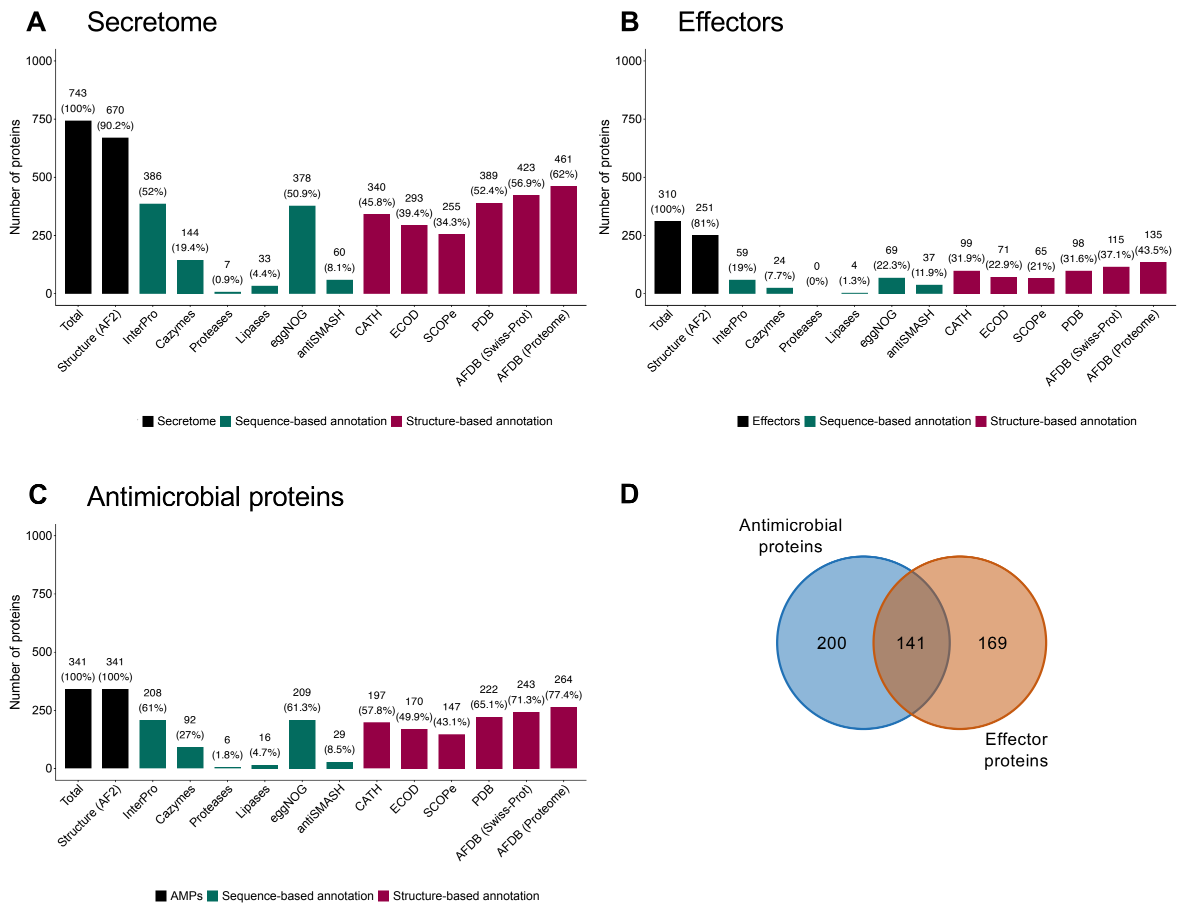


**Fig. S3. Summary of the number of proteins annotated across different databases.** Total numbers of proteins predicted and annotated using sequence-based and structure-based information are shown for **A)** Secreted, **B)** Effector, and **C)** Antimicrobial proteins. Predicted structures from AlphaFold2 (AF2) were used for structure-based annotation. Total predicted proteins and protein structures are shown in black, sequence-based annotations in teal, and structure-based annotations in magenta. The percentage of proteins annotated by each database refers to the total predicted in each category (secreted, effector, or antimicrobial proteins). **D)** Venn diagram comparing the number of secreted proteins predicted to have antimicrobial activity and proteins predicted as effectors. The overlap corresponds to 141 putative antimicrobial effectors.

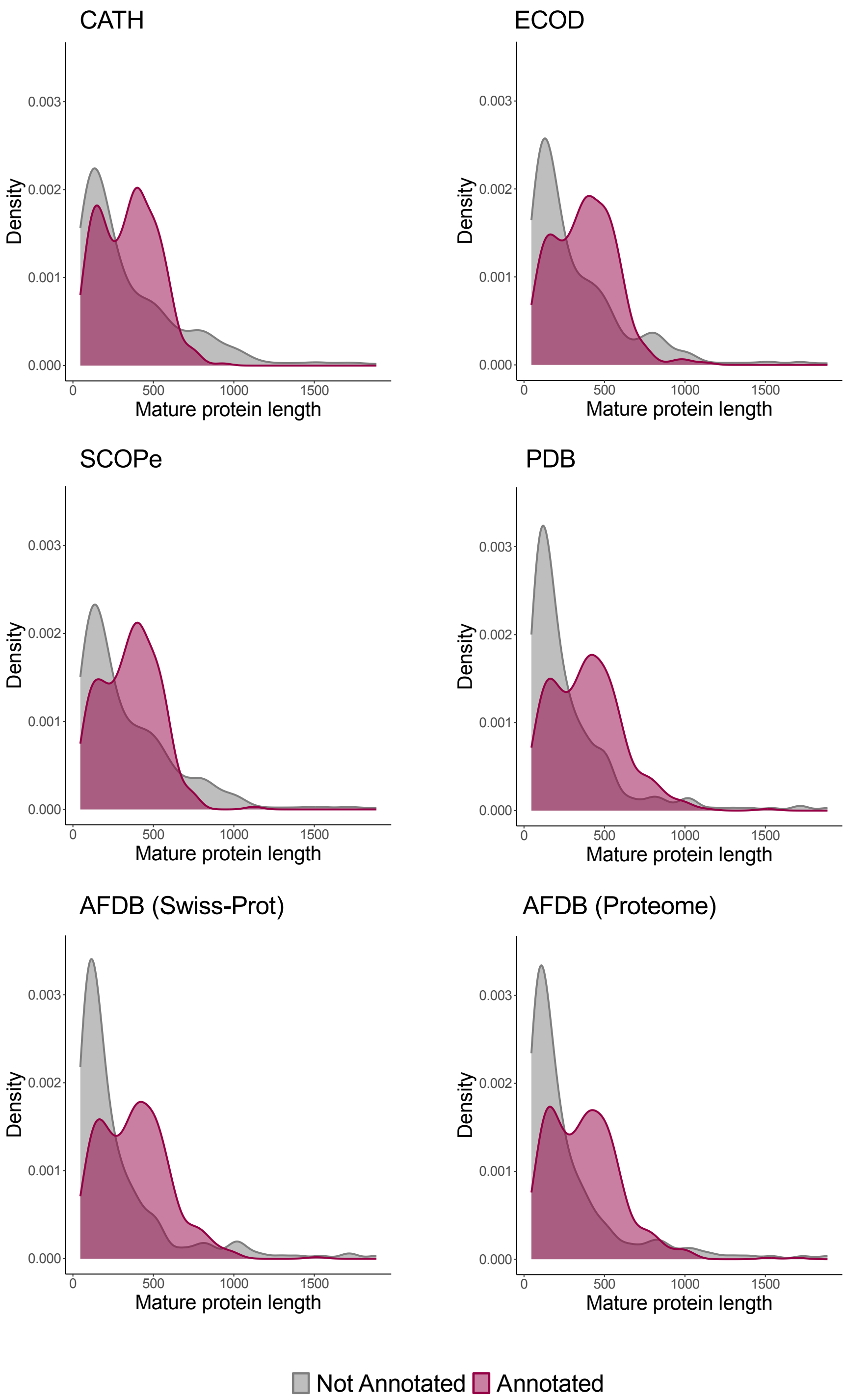


**Fig. S4. Distribution of annotated and non-annotated proteins across structural databases by mature protein length.** Each plot displays the probability density of proteins annotated (magenta) and non-annotated (grey) in each structural database.

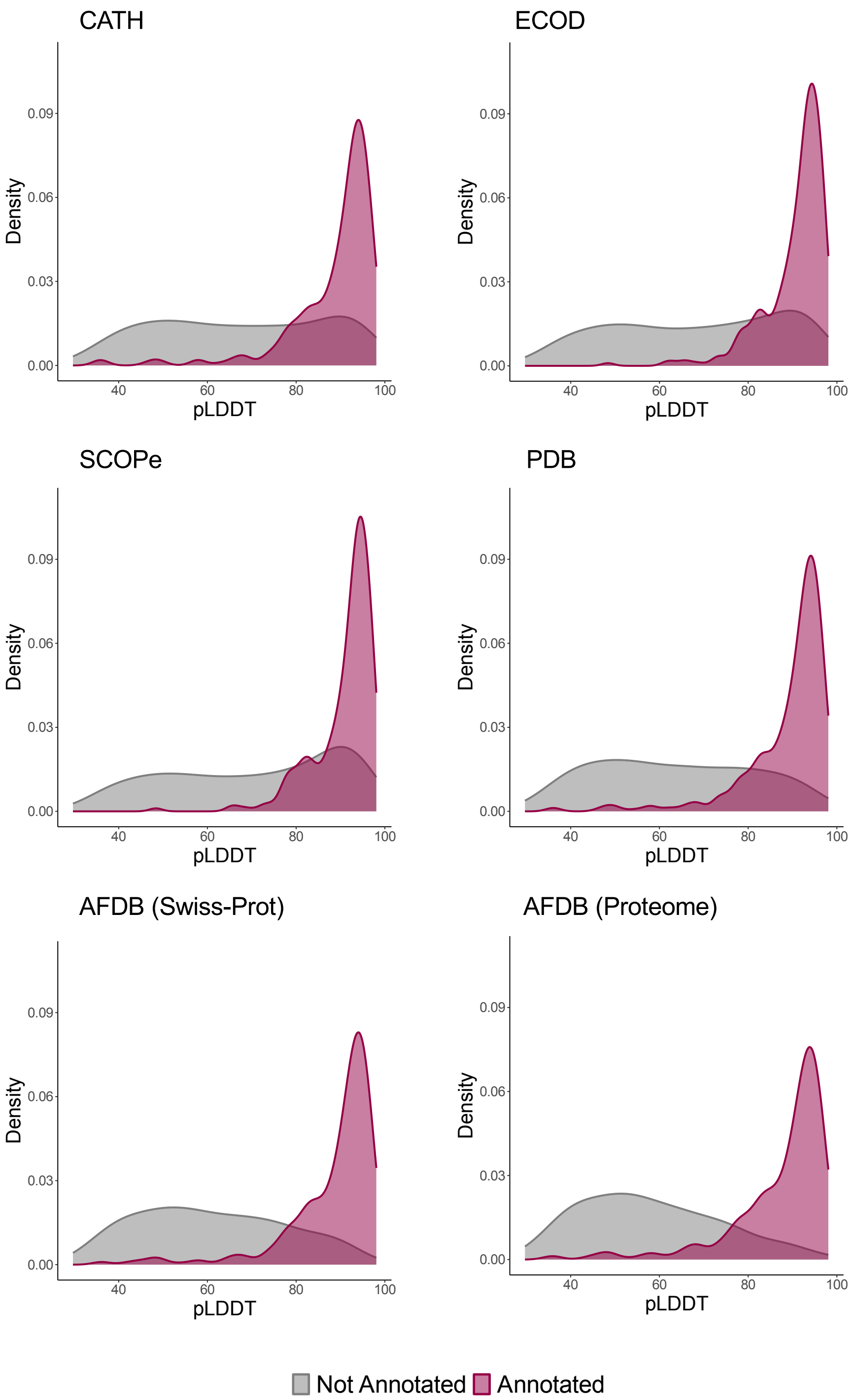


**Fig. S5. Distribution of annotated and non-annotated proteins across structural databases by pLDDT value.** Each plot displays the probability density of proteins annotated (magenta) and non-annotated (grey) in each structural database.

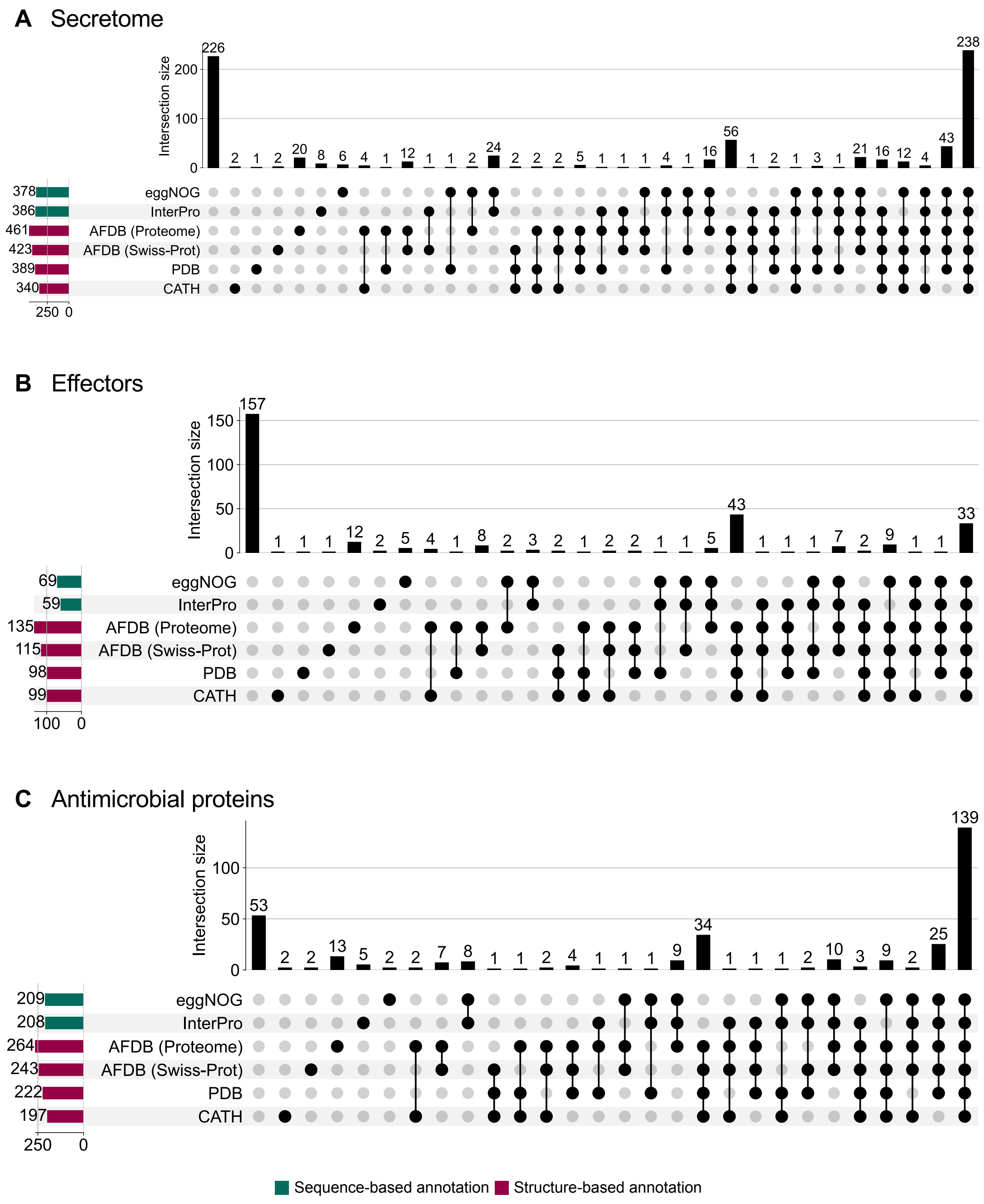


**Fig. S6. Overlap of protein annotation using sequence-based and structure-based information for A) Secreted, B) Effector, and C) Antimicrobial proteins**. Upset plots display the counts for the main databases shown in Fig. S3. Sequence-based annotations are shown in teal, and structure-based annotations in magenta.

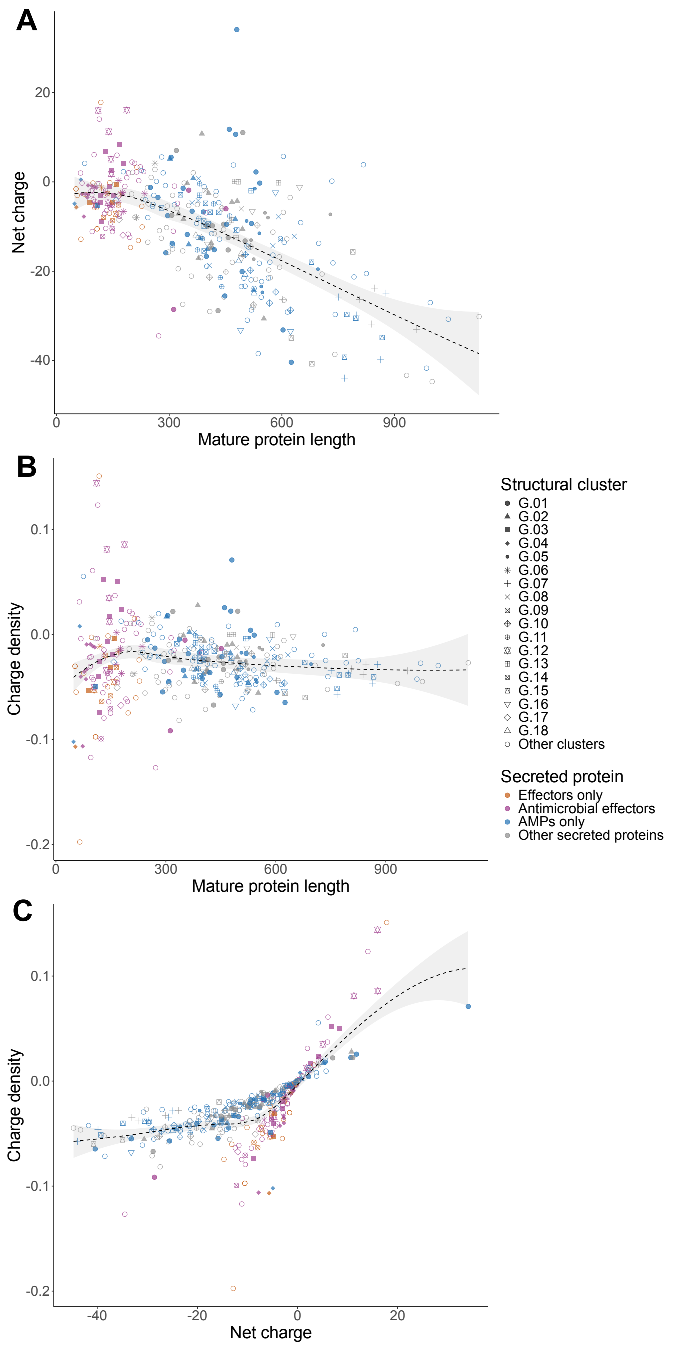


**Fig. S7. Pairwise distributions of net charge, absolute charge density, and mature protein length for proteins in the structural similarity network. A)** Net charge (pH 7) and mature protein length, **B)** Charge density (net charge per residue) and mature protein length**, C)** Charge density and net charge. Each point is a protein, colored according to the functional protein group (putative effectors only, putative antimicrobial effectors, predicted AMPs only, and other secreted proteins), and point shapes denote structural clusters. Dashed lines are LOESS fits with 95% confidence bands shaded in grey.

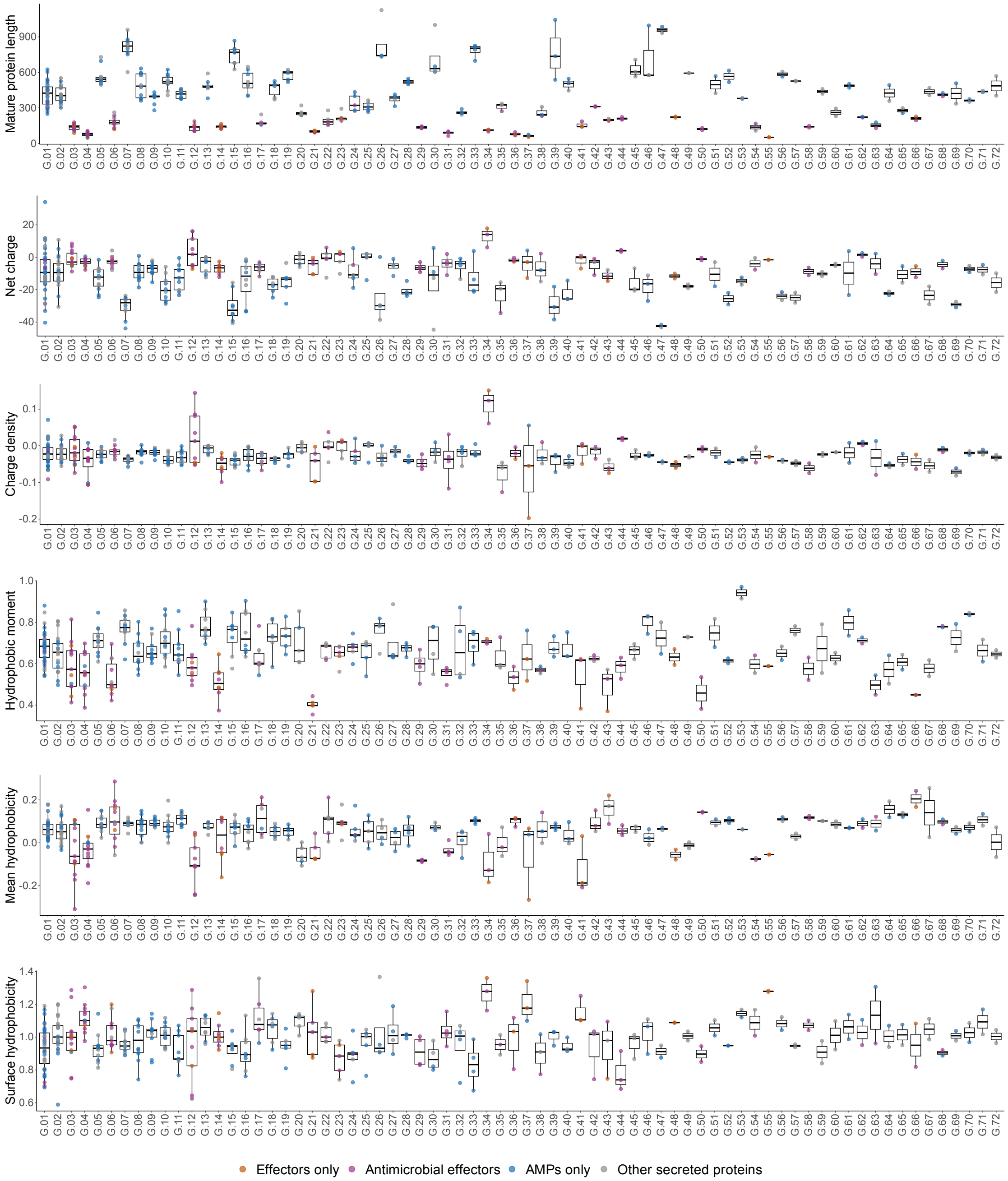


**Fig. S8. Distributions of six physicochemical properties across structural clusters of *Zymoseptoria passerinii* Zpa796.** From top to bottom, boxplots by cluster for: mature protein length, absolute net charge (pH 7), charge density (net charge per residue), hydrophobic moment (maximum local helical amphipathicity), mean hydrophobicity, and surface hydrophobicity. Each point is a protein, colored according to the functional protein group (putative effectors only, putative antimicrobial effectors, predicted AMPs only, and other secreted proteins). Pairwise differences were assessed among clusters G.01 to G.46, with at least three protein members; results according to Dunn's test are provided in Additional File 1: Table S11.

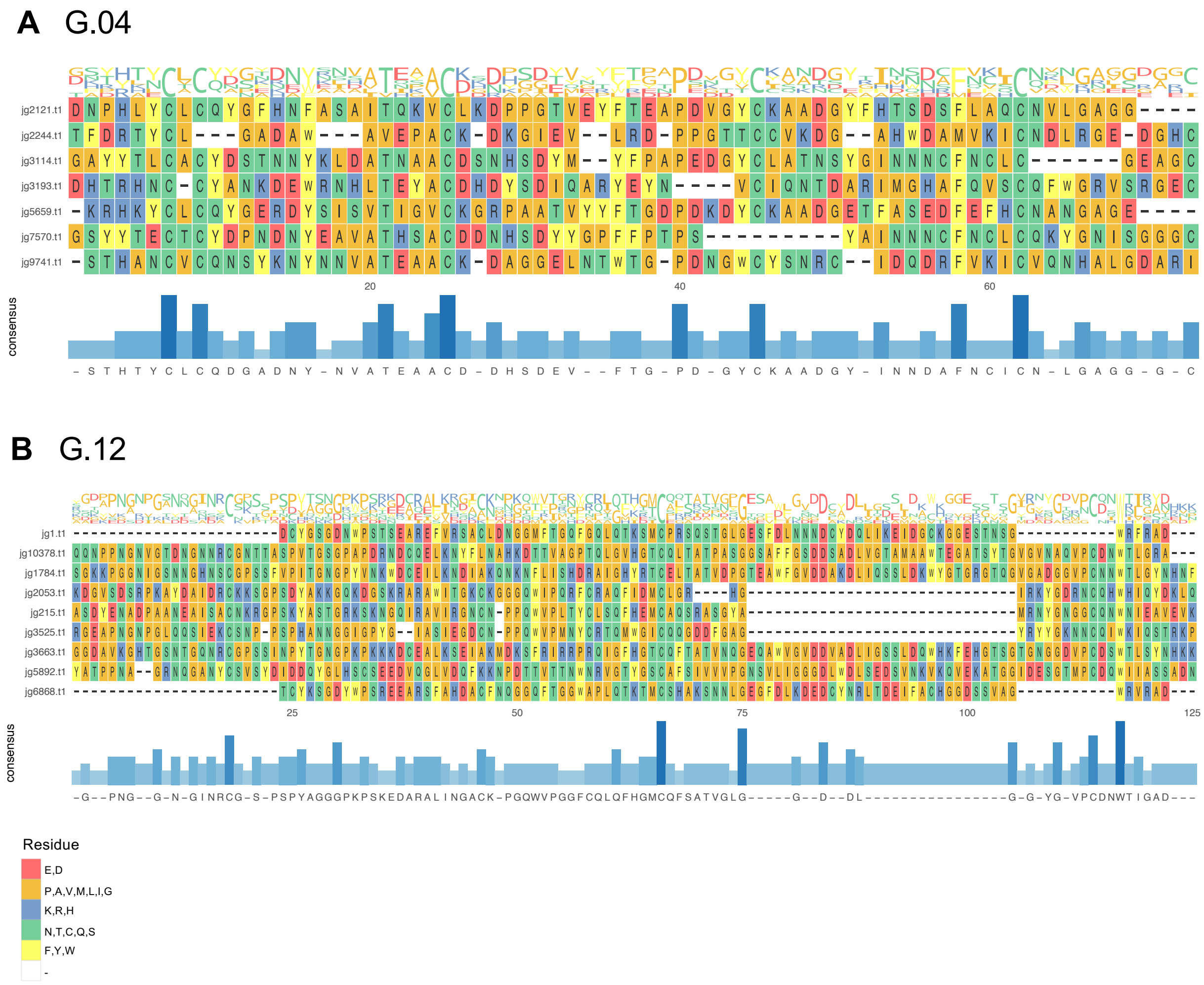


**Fig. S9. Trimmed multiple sequence alignment (MSA) of proteins from structural clusters A) G.04 and B) G.12.** Each alignment is presented in three panels: the upper panel shows the sequence logo, the middle panel displays the amino acid alignment, and the lower panel presents the consensus pattern of amino acid conservation. MSAs were generated using the L-INS-i algorithm from MAFFT and trimmed using the option --gappyout from trimAl.

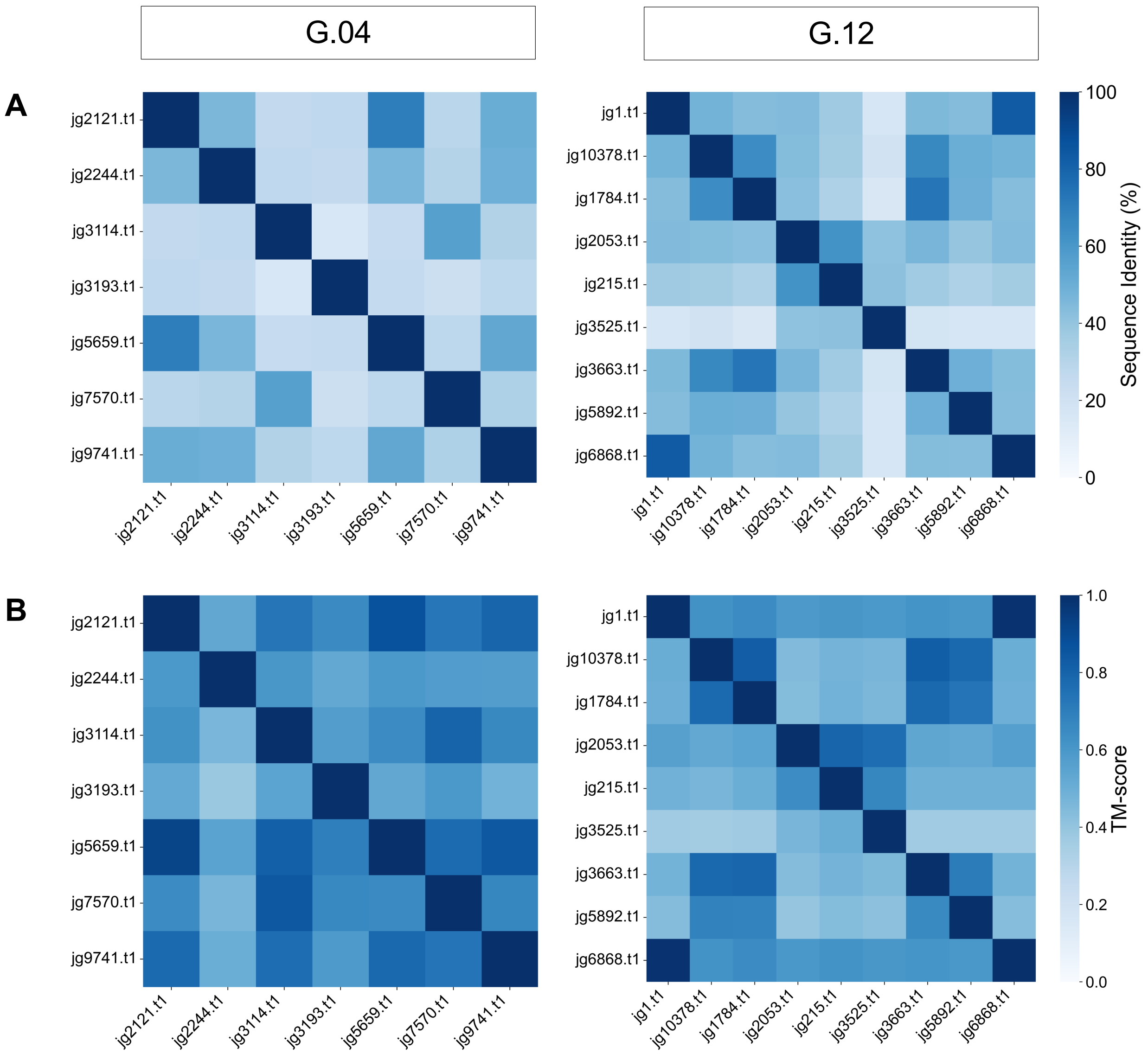


**Fig. S10. Pairwise comparison of sequence identity and structural similarity within clusters G.04 and G.12. A)** Percent sequence identity (symmetric matrix). **B)** TM-score (asymmetric matrix). In B, rows are query proteins and columns are subject proteins, with values normalized by the query length.

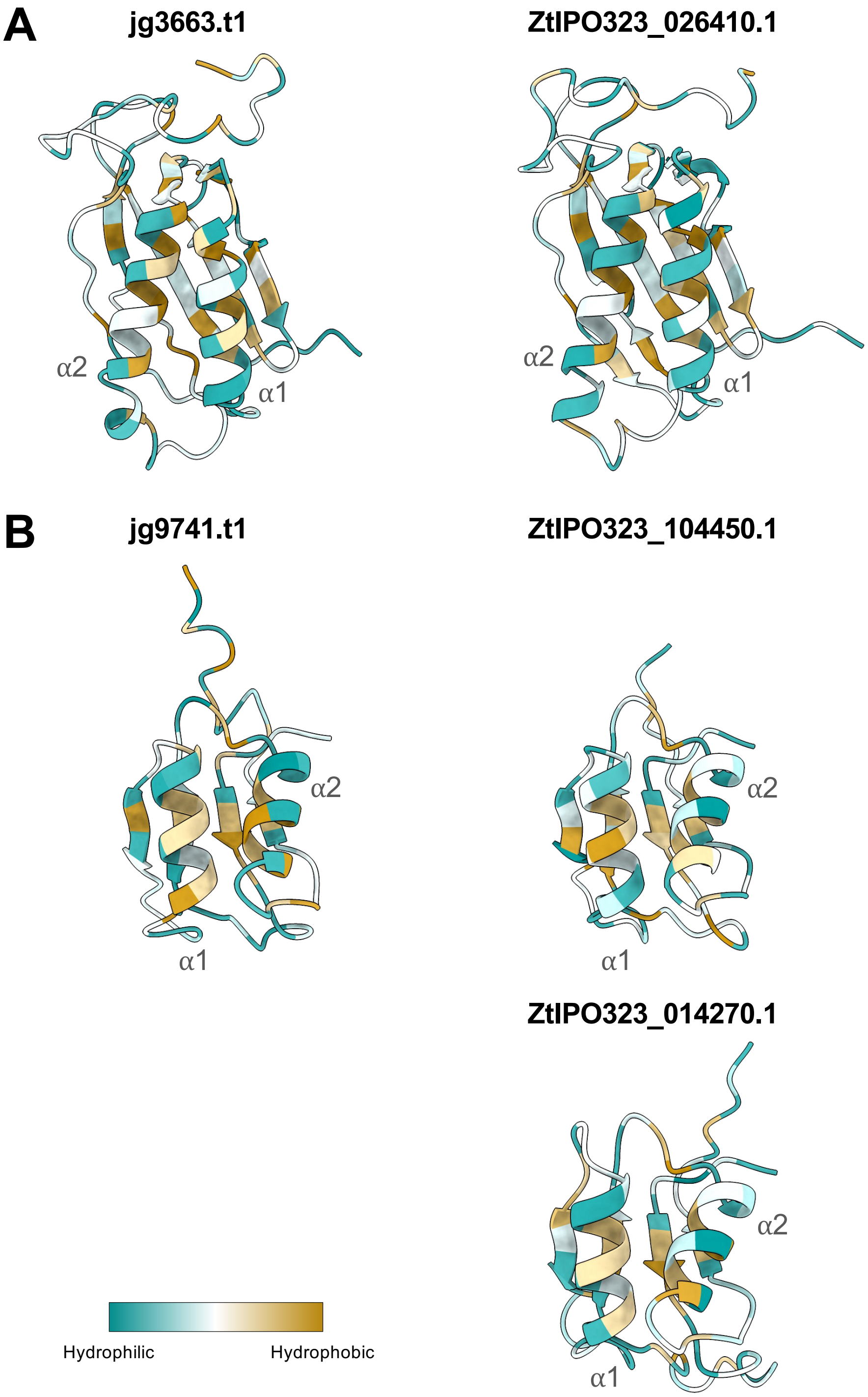


**Fig. S11. Comparison of the extent of hydrophobicity of *Zymoseptoria passerinii* Zpa796 proteins and their homologs in *Z. tritici* IPO323.** AlphaFold2 predicted protein structures are colored according to the Kyte–Doolittle hydrophobicity scale, with hydrophobic and hydrophilic regions mapped per residue.
