## Additional File 3 for "Conserved protein folds underpin the diversification of secreted proteins in a fungal pathogen"

### SUPPLEMENTARY RESULTS

#### Comparison of structural predictions from AlphaFold2 and ESMFold

To compare how similar AlphaFold2 (AF2) [1] and ESMFold [2] predictions are to experimentally validated fungal protein structures, we aligned crystallized fungal proteins retrieved from Protein Data Bank (PDB) [3] to their respective AF2- and EMSFold-predicted models (Additional File 2: Fig. S2). In total, we retrieved 228 crystallized protein structures and their amino acid sequences were used as input for AF2 and ESMFold calculations (Additional File 1: Table S3). AF2 successfully predicted structures for 226 proteins, but failed prediction for two small proteins from PDB, entries 2ALC (a DNA-binding protein from *Aspergillus nidulans*) and 2HGO (the cassiicolin toxin from the plant pathogen *Corynespora cassiicola*) (Additional File 1: Table S3). ESMFold successfully predicted structures for all 228 proteins.

When we compared the confidence of prediction methods, AF2-predicted structures exhibited significantly higher mean pLDDT values (93.36) than those predicted by ESMFold (80.82) for fungal proteins retrieved from the PDB ($p$-value $< 0.001$). Next, we assessed the structural similarity between AF2 and ESMFold models relative to crystallized structures from the PDB (Additional File 1: Fig. S2). AF2 predictions showed significantly higher mean template modeling (TM) score and lower root mean square deviation (RMSD) values when compared to PDB structures ($p$-value $< 0.01$) (Additional File 2: Fig. S2A-B).  We observed significantly higher TM-scores and lower RMSD values for PDB–AF2 alignments in proteins ranging from 101 to 700 amino acids long (Additional File 1: Fig. S2C-D), reflecting greater structural similarity between AF2 predictions and the crystalized fungal structures from PDB than ESMFold predictions compared to the same PDB proteins.

**Structural annotation of *Z. passerinii* Zpa796 secreted proteins**

We selected the 670 AF2-predicted structures of *Z. passerinii* Zpa796 for our downstream analyses (Fig. 2, Additional File 1: Table S4). A total of 479 (71.49%) protein structures from Zpa796 were successfully annotated, that is, they shared a similar fold with entries in at least one of the six structural databases we analyzed (Fig. 2, Additional File 1: Table S4). The highest number of protein structures shared a similar fold with entries in the AFDB [4] , with 461 and 423 annotated proteins with the AFDB (Proteome) and AFDB (Swiss-Prot) sections, respectively (Fig. 2A, Additional File 2: Figs. S3A and S6A). Zpa796 proteins also matched experimentally determined structures, with 389 proteins identified with a similar fold in PDB (Figs 2A, Additional File 2:  S3A and S6A). Structural databases (CATH, ECOD, and SCOPe) [5–7] had fewer matches to Zpa796 proteins (340, 293, and 255, respectively) (Fig. 2A, Additional File 2: Figs. S3A and S6A). We found overlapping for structures annotated across the databases, with 247 structures from Zpa796 sharing a similar fold entry in all six structural databases (Fig. 2A). Smaller overlaps for combinations of databases were also observed. In total, 46 proteins structures of Zpa796 had a best hit in only one of the six structural databases, with 38 of these found exclusively in the AFDB (Proteome). No exclusive best hits were found in the ECOD and SCOPe databases.

We explored the distribution of annotated structures according to mature protein length (Fig. 2B-C, Additional File 2: Fig. S4). Proteins ranging from 101 to 600 amino acids were annotated more frequently than shorter proteins (up to 100 residues) or longer ones (more than 600 residues) (Fig. 2B-C). The distribution of annotated proteins by length and structural confidence (pLDDT) followed similar trends across all databases, peaking at 401 amino acids and pLDDT = 94.26 (Fig. 2C-D, Additional File 2: Figs. S4-S5).

**Structure-based annotation of effector and antimicrobial proteins**

Using EffectorP [8] we predicted 310 effectors in the secretome of *Z. passerinii* Zpa796 (Additional File 2: Fig. S3B, Additional File 1: Table S1). We predicted structures for 251 (80.98%) putative effectors (Additional File 2: Fig. S3B) using AF2 and annotated 143 (56.97%) of them. Zpa796 putative effectors shared a similar fold with entries in AFDB (135 in AFDB (Proteome), 115 in AFDB (Swiss-Prot)), followed by entries in CATH (99 similar entries), PDB (98), ECOD (71), and SCOPe (65) (Additional File 2: Fig. S3B). In total, 78 putative effectors were exclusively annotated by at least one of the four main databases (AFDB (Proteome and Swiss-Prot sections), CATH, PDB) (Additional File 2: Fig. S6B).

Using AF2-predicted structures as input and the machine learning pipeline AMAPEC [9], we predicted 341 antimicrobial proteins (AMPs) (Additional File 2: Fig. S3C) and annotated 273 (80.06%) of them. The number of predicted AMPs that shared a similar fold in the structural databases were: 264 (AFDB (Proteome)), 243 (AFDB (Swiss-Prot)), 222 (PDB), 197 (CATH), 170 (ECOD), and 147 (SCOPe) (Additional File 2: Fig. S3C). In total, 68 predicted AMPs were exclusively annotated by at least one of the four main databases (AFDBs (Proteome and Swiss-Prot), CATH, PDB) (Additional File 2: Fig. S6C).

### Sequence-based annotation of the secretome

In addition to the structure-based annotation, we annotated the 743 predicted secreted proteins using amino acid sequences to identify conserved domains (Additional File 2: Fig. S3A, Additional File 1: Table S1). The sequence-based databases that annotated the highest number of secreted proteins were InterPro (386 annotated proteins, 51.95% of total secreted proteins) and eggNOG (378 proteins, 50.87%) (Additional File 2: Fig. S3A). When compared to the four main structural databases used (AFDBs – both Proteome and Swiss-Prot sections, CATH, PDB), only 38 proteins (5.11% of total secreted proteins) were annotated exclusively by InterPro and eggNOG (Additional File 2: Fig. S6A). Among the 310 putative effectors, InterPro and eggNOG annotated 59 (19.03%) and 69 (22.26%) proteins, respectively (Additional File 2: Fig. S3B). Only 10 (3.23%) effectors were annotated exclusively by one of these two sequence-based databases and not by any of the four main structural databases (Additional File 2: Fig. S6B). Among the 341 predicted AMPs, InterPro and eggNOG annotated 208 (61%) and 209 (61.30%) proteins, respectively (Additional File 2: Fig. S3C). Fifteen (4.40%) predicted AMPs were exclusively annotated by one of these two sequence-based databases when compared to structural annotations (Additional File 2: Fig. S6C). In total, 226 (30.42%) secreted proteins, 157 (50.65%) putative effectors, and 53 (15.54%) predicted AMPs were not annotated by any of the two sequence-based databases (InterPro, eggNOG) and the four structural-based databases (AFDB (Proteome), AFDB (Swiss-Prot), CATH, and PDB) (Additional File 2: Fig. S6).

#### Network statistics

To characterize the topology of the structural similarity network, we computed general network statistics and centrality measures across all nodes. Overall, network statistics were consistent with the fragmented nature of the network. The average degree was 4.813, indicating that each protein was structurally similar to roughly five others on average. The network density was 0.013, supporting a sparse network with few connections among all possible protein pairs. For centrality measures, the average degree centrality was 0.013, reflecting the proportion of nodes each protein is directly connected to. The average betweenness centrality was close to 0.000, indicating that most proteins do not act as bridges. The average closeness centrality was 0.016, supporting long paths between proteins within the network. Due to the high fragmentation of the network, we did not include statistics based solely on the largest connected component (i.e. average clustering coefficient, shortest path length, or network diameter) as these metrics would not capture the global architecture of the network.

#### Functional landscape of structurally similar clusters

Although not significantly enriched after p-value correction ($p\text{-value}_{adj} > 0.05$), 10 small structural clusters (composed of 2–4 proteins) contained only proteins predicted to be effectors (Fig 3). Some of these clusters were annotated into known protein folds or related to characterized biological functions, including: G.31 (OB-Fold; nucleotide-binding and antifungal activity), G.34 (Translation initiation factors EIF1), G.36 (Hydrophobins; structural proteins), G.41 (Pectate-lyases C), G.44 (Glucanases), and G.50 (Barwin-domain, endoglucanases) (Additional File 1: Table S4). Moreover, five small clusters

(G.29, G.31, G.44, G.50, and G.58) were composed exclusively of putative antimicrobial effectors (Fig. 3).

### Physicochemical properties of proteins across the network

To contextualize the lack of AMP enrichment in the network, we compiled six physicochemical properties and compared these properties across the structurally similar clusters (Additional File 2: Fig. S8, Additional File 1: Tables S9-S11). We limited comparisons to the 46 structural clusters with at least three proteins. The six properties showed significant differences across structural clusters according to Kruskal-Wallis tests (p-value < 0.001) (Additional File 1: Table S9). Post hoc pairwise comparisons using Dunn's test showed several significant differences ($p\text{-value}_{adj} < 0.05$) between clusters for five properties (Additional File 1: Table S11). There were 495 significant differences between pairs of structural clusters for mature protein length, 123 for net charge, 81 for hydrophobic moment, 75 for mean hydrophobicity, and one for surface hydrophobicity ($p\text{-value}_{adj} < 0.05$) (Additional File 1: Table S11). Overall, we identified significant differences between effector-enriched and effector-mainly clusters when compared to clusters composed of non-effector proteins (predicted as AMPs only and other secreted proteins) (Additional File 1: Table S11). Despite a significant Kruskal-Wallis test for charge density (Additional File 1: Tables S9), post hoc pairwise comparisons did not show any significant pairwise comparison between clusters after p-value correction ($p\text{-value}_{adj} > 0.05$) (Additional File 1: Table S11). This is likely because only the most pronounced cluster-specific differences were identified as statistically significant due to multiple-testing correction for 1035 comparisons using the Benjamini and Hochberg (BH) method.

### Comparison between homologs of *Z. passerinii* Zpa796 and *Z. tritici* IPO323

We used the OrthoFinder pipeline [10] to identify homologous relationships between proteins from *Z. passerinii* Zpa796 and other *Zymoseptoria* species. Among the 364 proteins from the structural similarity network of isolate Zpa796, five proteins — jg9754.t1 (G.03), jg9578.t1 (G.04), jg2591.t1 (G.07), jg2596.t1 (G.37), and jg5893.t1 (G.37) — had no homologs in any other *Zymoseptori*a species or in *Z. passerinii* Zpa63 (Additional File 1: Table S12), suggesting they may be isolate-specific. Among the remaining 359 Zpa796 proteins that shared homologs with at least one other species (Additional File 1: Table S12). Some had a single ortholog in all compared individuals, including the reference isolate *Z. tritici* IPO323 [11–13], such as the putative antimicrobial jg3663.t1 (G.12) (Additional File 1: Table S12). Others had variable number of homologs in the other species, such as jg9741.t1 (G.14). Homologs of this putative antimicrobial effector varied from one (in Z. *passerinii* Zpa63, *Z. pseudotritici* Zp13, and *Z. tritici* Zt469) [14,15] to five copies in *Z. ardabilae* Za17 [14] (Additional File 1: Table S12).

The structural alignments of *Z. passerinii* Zpa796 proteins (jg3663.t1 and jg9741.t1) and their homologs in *Z. tritici* IPO323 showed overall structural conservation, but we identified local variations primarily in loop regions (Fig. 6). The superposition of jg3663.t1 (G.12, KP4-like) and ZtIPO323_026410.1 structures showed a conserved core fold (TM-score = 0.87, RMSD = 2.08), with major differences in loop extensions (small disruptions reflecting indels in the diagonal pattern of the contact map, Fig. 6B) and terminal regions. Similarly, there was overall structural conservation between jg9741.t1 (G.04, alpha-beta plait/ferredoxin-like) and its IPO323 homologs ZtIPO323_104450.1 and

ZtIPO323_014270.1 (TM-scores ranging from 0.80 to 0.82 and RMSD values between 1.46 and 1.55) (Fig. 6A). The core helices and beta-strands were conserved, whereas we identified structural differences primarily in the loop regions, resulting from indels. The most prominent structural deviations were in the loop regions of the alignment between jg9741.t1 and ZtIPO323_014270.1, with the *Z. tritici* IPO323 protein having more extended and less constrained β1-α1 and β2-β3 loops than its homolog in *Z. passerinii* Zpa796 (Fig. 6).

### SUPPLEMENTARY METHODS

#### Sequence-based protein functional annotation

Protein domains were annotated using InterProScan v5.54-87.0 [16] with the applications: SMART-7.1, SUPERFAMILY-1.75, CDD-3.18, TIGRFAM-15.0, Pfam v34.0, and Gene3D-4.3.0. EuKaryotic Orthologous Groups (KOG), Gene Ontology, and KEGG terms were assigned to the proteome using eggNOG-mapper v2 [17] against the eggNOG 5.0 database [18]. Carbohydrate-active enzymes (CAZymes) were predicted using the dbCAN2 HMM profile database v7.0 [19] and *hmmscan* from HMMER v3.1b2 [20]. Lipases were predicted using HMM profiles from the Lipase Engineering Database v3.0 [21]and *hmmscan*. Significant matches for CAZymes and lipases were parsed using as cutoff E-value = 1e-4 and coverage = 0.35. Proteases were predicted using the fungal proteome as queries and the MEROPS database v12.1 [22] with BLASTp run locally (thresholds set as E-value = 1e-4, minimum identity = 0.40, coverage = 0.80). The secondary metabolite biosynthesis gene clusters in the genome were predicted with antiSMASH v7.1.0 [23] for fungi using the following features: KnownClusterBlast, ClusterBlast, SubClusterBlast, MIBiG Cluster, ActiveSiteFinder, RREFinder, Cluster Pfam, Pfam-based GO Term Annotation, and TIGRFam.

#### Fungal secretome modelling by AlphaFold2 and ESMFold

We compared structural prediction methods using the entire secretome of *Z. passerinii* Zpa796 and by sequence ranges based on the mature protein lengths. Proteins were grouped into 100-amino-acid length groups (e.g., 1–100, 101–200, up to proteins with more than 1,000 residues). To compare the quality of structural predictions by AF2 and ESMFold for secreted proteins, we examined the correlation between the two methods by comparing pLDDT values. We assessed data normality with the Shapiro-Wilk test (p-value $< 0.05$), followed by Spearman's correlation test. Distributions of pLDDT values for the entire predicted secretome and for each length category were compared using the Wilcoxon rank-sum test (with Benjamini and Hochberg (BH) adjusted p-values) and effect sizes were calculated using Cohen's d. Additionally, we used TM-align v20220412 [24] with default parameters to compare structural similarity between prediction methods by assessing the template modeling (TM) score and root mean square deviation (RMSD) values obtained from pairwise structural alignments of proteins with predicted structures by both methods. Differences in structural similarity across protein length categories were evaluated for each similarity index using the Kruskal-Wallis test followed by Dunn's test ($\text{p-value}_{adj} < 0.05$, BH correction). To measure how similar the AF2 and ESMFold predictions were to crystallized fungal protein structures, we compared

TM-scores and RMSD values between structures predicted by both methods and crystalized structures retrieved from the Protein Data Bank (PDB). Statistical comparisons were performed using the Kruskal-Wallis and Dunn's tests in R v4.3.3 [25].

### Physicochemical properties calculated by AMAPEC

Among the six physicochemical properties we compared for the secreted proteins of *Z. passerinii* Zpa796, four were already calculated as part of AMAPEC v1.0b pipeline [9]: net charge, mean hydrophobicity, and hydrophobic moment, and surface hydrophobicity. Net charge, mean hydrophobicity, and hydrophobic moment were computed from mature protein sequences using the *Peptides* package [26] in R v4.3.1, as implemented in AMAPEC with default parameters. Net charge was calculated at pH = 7 and using 'pKscale = "EMBOSS"'. Mean hydrophobicity was calculated using the Eisenberg scale and the arithmetic mean per-residue values across the mature protein sequence. Hydrophobic moment (amphipathicity) was quantified with a protein rotational angle of 100 and a sliding window of 11 residues; for each protein, the maximum value across all sliding windows was retained to capture the strongest local helical amphipathicity. Surface hydrophobicity was computed from solvent-accessible surface areas via Bio.PDB.DSSP module from Biopython v1.83 [27].
